# Optics-free reconstruction of shapes, images and volumes with DNA barcode proximity graphs

**DOI:** 10.1101/2024.08.06.606834

**Authors:** Hanna Liao, Sanjay Kottapalli, Yuqi Huang, Matthew Chaw, Jase Gehring, Olivia Waltner, Melissa Phung-Rojas, Riza M. Daza, Frederick A. Matsen, Cole Trapnell, Jay Shendure, Sanjay Srivatsan

## Abstract

Spatial genomics technologies include imaging- and sequencing-based methods. Sequencing-based spatial methods typically require surfaces coated with coordinate-associated DNA barcodes, but the physical registration of these barcodes to spatial coordinates is challenging, necessitating either high density printing of oligonucleotides or *in situ* sequencing/probing of randomly deposited, DNA-barcode-bearing beads. As a consequence, the surface areas available to sequencing-based spatial genomic methods are constrained by the time, labor, cost and instrumentation required to either print or decode a coordinate-tagged surface. To address this challenge, we developed SCOPE (Spatial reConstruction via Oligonucleotide Proximity Encoding), an optics-free, DNA microscopy-inspired method. With SCOPE, the relative positions of DNA-barcoded beads within a 2D shape, 2D image or 3D volume are inferred from the *ex situ* sequencing of chimeric molecules formed from diffusing “sender” and tethered “receiver” oligonucleotides. To demonstrate the potential of this approach, we applied SCOPE to reconstruct 2D shapes, 2D images or 3D volumes defined by 10^4^-10^6^ *x* 20-100 µm DNA barcoded beads, including an asymmetric “swoosh” resembling the Nike logo (44 mm^2^), a “color” Snellen eye chart (704 mm^2^) and the surface topology of 3D molds of a teddy bear, star, butterfly or block letter (75-100 mm^3^). Each of the resulting “DNA barcode proximity graphs” was computationally reconstructed in an automated fashion, across fields of view and at resolutions that were determined by sequencing depth, bead size and diffusion kinetics, rather than by microarray or microscope instrument time. Because the ground truth shapes are known, these datasets may be particularly useful for the further development of computational algorithms by this nascent field.

## INTRODUCTION

In the natural and engineered worlds, conventional paradigms for determining spatial relationships include optics (*e.g.* microscopy), acoustics (*e.g.* echolocation) and haptics (*e.g.* touch). However, particularly at nanoscopic and microscopic scales, one can also imagine how complex polymersーand DNA in particularー might be leveraged to encode and decode spatial information. For such a goal, DNA has a number of attractive properties, including: its high information density; simple design principles; biochemical stability; the breadth of primitive operations that can be accessed enzymatically or directly encoded (Kishi et al., 2018); and our contemporary access to low-cost technologies for synthesizing and sequencing nucleic acids of arbitrary sequence composition. Many of these same properties underlie DNA’s promise as a digital data storage medium, whether *in vitro* (*e.g.* using A’s, G’s, C’s and T’s to encode written symbols or visual pixels (Church et al., 2012; Davis, 1996)) or *in vivo* (*e.g.* using prime editing-mediated short insertions to encode ordered molecular signaling or cell lineage histories (Chen et al., 2024; Choi et al., 2022)).

DNA-based molecular barcoding (aka tagging or multiplexing) (Church and Kieffer-Higgins, 1988) is at the core of many modern genomic methods, including: (i) unique molecular identifiers (UMI), with which RNA molecules are tagged; (ii) massively parallel reporter assays (MPRAs), in which regulatory elements are tagged; (iii) pooled genetic screens, in which perturbations, variants or lineages are tagged; (iv) single cell combinatorial indexing, in which cells are tagged; (v) proximity ligation assays, in which proteins or nucleic acids are tagged; (vi) and spatial genomics, in which hybridization probes or spatial coordinates are tagged. In each of these methods, the barcode sequences themselves are usually arbitrary, while for some further applications (*e.g.* DNA origami (Rothemund, 2006), DNA computing (Benenson et al., 2001)), the barcode sequences and their interactions encode some pre-specified molecular logic.

Over the past decade, spatial genomic technologies have rapidly proliferated (Moses and Pachter, 2022; Vandereyken et al., 2023; Williams et al., 2022). The underlying methods can be dichotomized into those that rely on *in situ* imaging (Chen et al., 2015; Lubeck and Cai, 2012) vs. *ex situ* sequencing for primary data collection, while *ex situ* sequencing methods can be further dichotomized into those that rely on microdissection (Nichterwitz et al., 2016) vs. DNA-based molecular barcodes to identify the spatial coordinates associated with each sequencing read, such as Slide-seq (Rodriques et al., 2019), HDST (Vickovic et al., 2019), and DBiT-seq (Liu et al., 2020). The main advantages of *ex situ* sequencing methods include that they do not require *a priori* specification of molecular species of interest and are less dependent on the cost and throughput limitations of optical instrumentation, while a further advantage of the spatial barcoding subset of *ex situ* sequencing methods is that substantially greater resolution can be achieved than with microdissection.

However, a major limitation of spatial barcoding methods is that they require the deposition and mapping of specific barcoded oligonucleotides (oligos) to specific locations on a 2D surface. The mapping between specific barcodes and specific physical coordinates is analogous to the mapping between the elements of a CCD array and individual pixels in a digital camera. To fabricate and map arrays of DNA barcodes, oligos can be: (i) pre-synthesized, arrayed, and deposited to specific locations, *e.g.* by a microarray printer (Srivatsan et al., 2021; Ståhl et al., 2016); (ii) synthesized to specific locations *in situ (Kishi et al., 2022)*; or (iii) randomly distributed and decoded via *in situ* hybridization or sequencing (Chen et al., 2022; Fu et al., 2022; Rodriques et al., 2019). However, these approaches are all time-, cost-, labor- or instrumentation-intensive, which limits the physical dimensions of the arrays that are typically produced and used for these assays.

Around 2019 to 2020, several groups proposed that diffusion-driven interactions between closely located, barcoded molecules could reveal spatial relationships, because physically proximate barcodes would interact far more often than distant ones (Boulgakov et al., 2020; Hoffecker et al., 2019). These ideas build on the logic of proximity ligation assays (Fredriksson et al., 2002), which use barcodes to detect pairwise molecular contacts, but extend it to reconstructing the relative positions of entire populations of barcoded molecules based solely on interaction frequencies. Around the same time, Weinstein & colleagues reduced similar ideas to practice with “DNA microscopy”, a method in which endogenous transcripts in fixed cells or tissues are tagged with UMIs, amplified, concatenated and sequenced (Weinstein et al., 2019). As the probability of concatenation is a function of molecular proximity, the physical relationships among the original transcripts could be inferred from sequencing data with cellular resolution. Since 2020 and particularly in the last year, there has been growing interest in advancing the theoretical (Greenstreet et al., 2023), computational (Fernandez Bonet and Hoffecker, 2023; Kloosterman et al., 2024) and experimental (Abdulraouf et al., 2024; Hu et al., 2025; Karlsson et al., 2024; Qian and Weinstein, 2025) frameworks underpinning this nascent class of methods.

Here we describe SCOPE (<u>S</u>patial re<u>C</u>onstruction via <u>O</u>ligonucleotide <u>P</u>roximity <u>E</u>ncoding), a DNA microscopy-inspired, optics-free method in which a 2D array or 3D volume of randomly deposited beads, each bearing a unique DNA barcode, self-register their relative positions via proximity-dependent hybridization and overlap extension of released “sender” and tethered “receiver” oligos. Massively parallel sequencing of sender-receiver chimeras results in a matrix of pairwise counts. Particularly for local communities, applying dimensionality reduction algorithms widely used for scRNA-seq analysis, such as Uniform Manifold Approximation and Projection (UMAP), to these data results in reasonable reconstructions of the relative positions of beads underlying each shape, image or volume (McInnes et al., 2018; van der Maaten and Hinton, 2008). As a first proof-of-concept (<u>2D shape</u>), we apply SCOPE to reconstruct an asymmetric “swoosh” resembling the Nike logo. As a second proof-of-concept (<u>2D image</u>), we use oligos and a microarray printer to encode a multi-color image of the Snellen eye chart for visual acuity, which we then reconstruct with SCOPE. As a third proof-of-concept (<u>3D volume</u>), we apply SCOPE to reconstruct the surface topology of 3D molds of a teddy bear, star, butterfly and block letter. Collectively, these demonstrations—spanning 10⁴–10⁶ barcoded beads (20–100 µm)—showcase the potential of DNA barcode proximity graphs for optics-free encoding and recovery of spatial information from 2D arrays or 3D volumes of beads. Given that the original 2D and 3D shapes or images are known, they may also provide “gold standard” datasets for developing and benchmarking computational algorithms for optics-free reconstruction.

## RESULTS

### Overview of SCOPE

We set out to develop a method for generating arbitrarily large surfaces or volumes of DNA barcodes whose relative spatial positions could be determined without relying on optical or microjet instrumentation. The strategy that we devised is based on a population of beads bearing unique DNA barcodes (*i.e.* wherein any given bead bears many copies of a single DNA barcode, while different beads bear distinct DNA barcodes). However, the oligos tethered to any given bead are functionally heterogeneous: they include both “messengers”, designed to interact with messengers from nearby beads, and “decoders”, designed to capture molecules of interest (**Fig. 1A**). Furthermore, each of these categories include further subsets termed “senders”, designed for programmed release, and “receivers”, which remain tethered to a given bead (**Fig. 1A**). After generating a 2D array or 3D volume of densely packed, DNA barcode-bearing beads, sender-messengers are released to diffuse and hybridize to receiver-messengers that remain tethered to proximally located beads (**Fig. 1B**). Massively parallel sequencing of sender-receiver messenger chimeras results in a matrix of barcode-barcode interactions that is informative with respect to the proximity relationships of beads, enabling computational reconstruction of their spatial arrangement (**Fig. 1C**). In parallel, decoder-receivers can capture externally sourced, proximal nucleic acids, enabling spatial mapping of additional molecular information onto the reconstructed map of bead positions (**Fig. 1B-C**).

**Figure 1:**
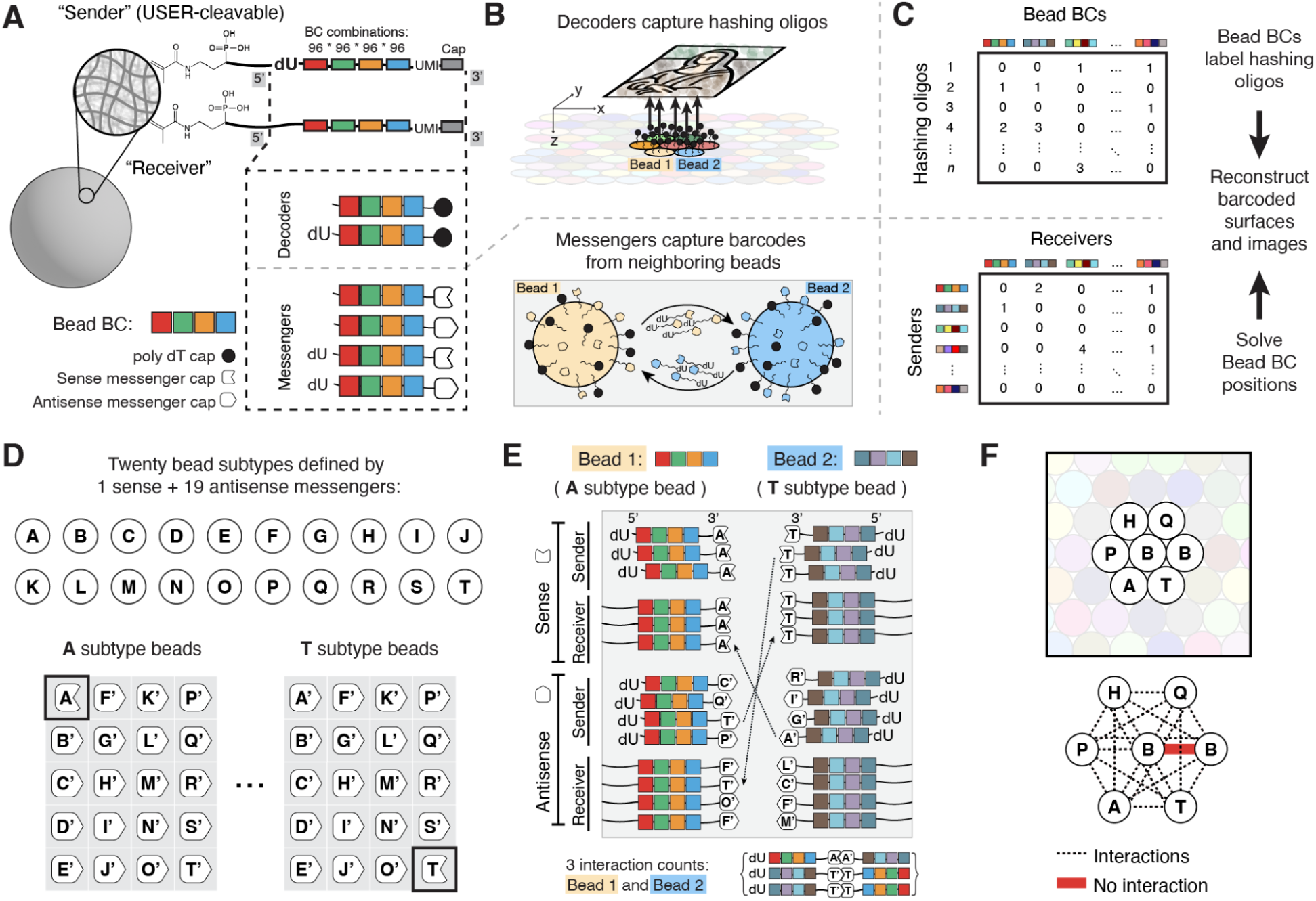
Overview of SCOPE (<u>S</u>patial re<u>C</u>onstruction via Oligonucleotide Proximity Encoding). **(A)** Schematic of structure of oligos tethered to SCOPE beads. Hydrogel beads contain 5’-Acrydite-tethered DNA oligos that are either cleavable (“senders”) or non-cleavable (“receivers”) by USER enzyme, based on the presence or absence of dU in the 5’ stem. Both sender and receiver oligos contain the same barcode structureー4 subsequences each with 96 possible identitiesーdepicted by the 4 colored boxes. Both sender and receiver oligos are also functionalized with a 3’ cap of either poly-dT (“decoders”, designed to capture poly-dA or poly-A tailed molecules) or a sequence encoding bead subtype (“messengers”, designed to chimerize with messenger oligos from proximally located beads). **(B)** Schematic of 2D SCOPE reaction. Top: Poly-dA hashing oligos are deposited onto a 2D array of SCOPE beads, bearing subsequences as “colors” that encode an image of interest. Decoder oligos form chimeras with proximally located hashing oligos. Bottom: Messenger oligos participate in diffusion-dependent, hybridization-mediated reactions with the messenger oligos of proximally located beads in the 2D array, resulting in sender-receiver messenger chimeras. **(C)** Top: Spatial mapping of hashing oligos. From sequencing of decoder-hashing oligo chimeras, we obtain a bead-by-molecule count matrix that is informative with respect to which hashing oligos are proximate to each bead barcode. Bottom: Inference of the relative positions of beads. From sequencing of messenger chimeras, we obtain a bead-by-bead count matrix that is informative with respect to which bead barcodes are proximate to which other bead barcodes. **(D)** Out of 20 designed pairs of complementary messenger sequences, the messenger oligos of each bead subtype are capped with 19 “antisense” sequences and 1 “sense” sequence. For example, the messenger oligos of “A” subtype beads include the “A” hybridization sequence in sense orientation, with the other 19 hybridization sequences in antisense orientation. **(E)** Illustration of how 3 interaction counts between a proximally located pair of “A” and “T” subtype beads might originate from different kinds of sender-reciever chimeras. **(F)** Only beads of different subtypes can contribute to the generation of sender-receiver chimeras, which avoids “self-reactions” (*i.e.* senders chimerizing with receivers from the same bead) but runs the risk of missing interactions between closely located beads of the same subtype.

### Generation of SCOPE beads

We chose to generate 20 µm hydrogel beads as a solid substrate for SCOPE’s proof-of-concept, because low-cost, accessible methods exist for the combinatorial DNA barcoding of such beads (Delley and Abate, 2021), as well as for generating densely packed, large 2D arrays from them. Additionally, acrylamide hydrogels have a high internal porosity which facilitates a higher density loading of oligos compared to non-porous substrates like glass or polystyrene (Fu et al., 2022). We generated 20 µm polyacrylamide hydrogel beads containing 100 µM of 5’-Acrydite oligo using a flow-focusing microfluidics chip (**Fig. S1A**). To facilitate the enzymatic release of sender molecules, half of the 5’-Acrydite oligos were synthesized containing a deoxyuracil (dU) in the 5’ stem (**Fig. 1A**). Subsequent split-pool combinatorial DNA synthesis via programmed ligation generated a combinatorial space of approximately 85 million possible barcodes comprising four positions with 96 possible sub-barcodes at each (Delley and Abate, 2021). Sequencing of the synthesized barcodes confirmed that the process was highly efficient, with sub-barcodes that were uniformly incorporated at each round (**Fig. S1B-C**). After barcoding, the bead-tethered oligos were then 3’ capped with sequences enabling either decoder or messenger functionalities (**Fig. 1A**). Through this process, the oligos decorating any single bead are homogenous in that they bear the same barcode, but heterogeneous in that sender-decoders, receiver-decoders, sender-messengers, and receiver-messengers, are all represented among them (**Fig. 1A**). Based on amplification and sequencing of UMIs associated with barcodes enzymatically released from individual beads, we estimate that each bead bears ∼500,000 functionalized, barcoded oligos (mean: 491,811; IQR: 308,757-833,744; **Fig. S1D**). To generate a planar array, SCOPE beads can be cast within an encasing hydrogel that polymerizes directly onto a treated glass slide (**Fig. S1E-F**).

### Messenger subtypes minimize the likelihood of self-self chimeras

SCOPE requires that sender and receiver messengers have some region of complementarity to hybridize to one another. This scheme bears some risk of “self-self” interactions, or chimeras between senders and receiver messengers derived from the same bead, a challenge that also plagues other proximity-dependent molecular methods such as Hi-C (Lajoie et al., 2015). To minimize this risk, we designed 20 distinct “cap” species for messengers (**Fig. 1A**; **Fig. S2A**). After synthesizing DNA barcodes but prior to capping, beads were split into 20 subsets and loaded with a mixture of messenger cap species, such that DNA barcodes of each “bead subtype” were capped with the antisense sequence of 19 cap designs, and the sense sequence of the remaining cap design (**Fig. 1D**). As a consequence, the messengers associated with a given bead subtype (each named for its sense-oriented cap) are capable of hybridizing with the messengers of all 19 other bead subtypes, but not messengers of its own subtype (**Fig. 1E**). This scheme avoids “self-reactions” (*i.e.* senders chimerizing with receivers derived from the same bead), but runs the risk of missing interactions between neighboring beads of the same subtype (**Fig. 1E-F**). The choice of 20 bead subtypes was based on simulations indicating diminishing returns beyond that point for minimizing the risk of the latter scenario (**Fig. S2B**).

### SCOPE reaction and massively parallel sequencing of sender-receiver chimeras

The SCOPE reaction involves subjecting the hydrogel-encased monolayer of polyacrylamide beads (**Fig. S1E-F**) to a temperature-controlled, two-phase reaction mediated by a single reaction volume containing two enzymes. In the first phase (37°C; 15’), sender-messengers are released from beads by the USER enzyme, which cleaves at the dU present at the stem of a subset of DNA barcodes (**Fig. 1A**). Sender-messengers diffuse away and hybridize to receiver-messengers tethered to adjacent beads (**Fig. 1B,E-F**). Because the complementary regions reside in the functional cap (**Fig. 1A**), senders and receivers are expected to hybridize at their 3’ ends. In the second phase (55°C; 15’), reverse transcriptase, which can mediate both RNA- and DNA-templated DNA polymerization (Hu and Hughes, 2012), drives overlap extension and the generation of a chimeric molecule that includes both the sender-messenger and receiver-messenger derived DNA barcodes (or, alternatively, chimeras of decoders and proximate hashing oligos) (**Fig. 1B**). Sender-receiver chimeras are then PCR amplified and subjected to massively parallel sequencing, which yields a sparse matrix of interaction counts between a large number of DNA barcode-defined beads. As the SCOPE reaction is dependent on a sender-messenger diffusing into the vicinity of a receiver-messenger, we expect the count frequencies between any given pair of beads to be a function of their physical distance. Importantly, the asymmetry of primer binding sites is such that for any given chimera, we are aware of which participating bead contributed the sender-messenger vs. the receiver-messenger. The resulting matrix of count frequencies is therefore asymmetric (**Fig. 1C**).

### SCOPE results in count matrices that are spatially informative

With hydrogel beads, we can readily generate 2D arrays of beads with a mean diameter matching expectation (20 µm) and 76% (s.d. 5%) coverage of the total area (**Fig. 2A,C**). As the upper bound on coverage is 91% (perfectly uniform, hexagonally packed circles), our packing could potentially be improved by improving bead size uniformity. With hexagonal packing, each bead has six immediate neighbors. Using image segmentation and Delaunay triangulation on a representative image of a SCOPE bead array, we found the mode number of immediate neighbors of a bead to be six (**Fig. 2B,D**).

**Figure 2:**
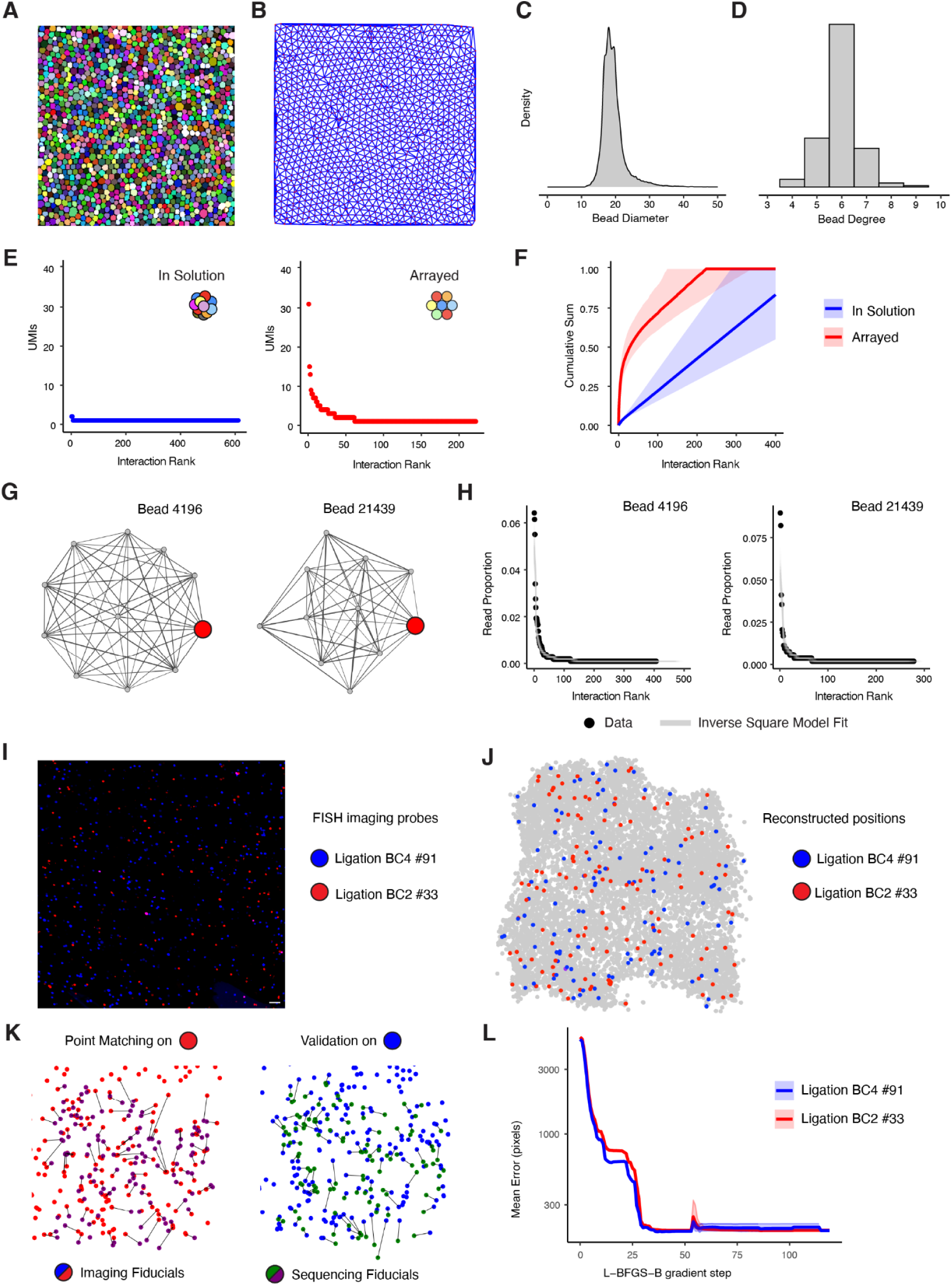
Bead arrays form local communities that can be computationally reconstructed from pairwise interaction matrices. **(A)** Segmented mask of a representative image of a SCOPE bead array. **(B)** Delaunay triangulation for segmented bead centroids. **(C)** Density plot of bead diameters from the segmented image. **(D)** Distribution of the number of nearest neighbors observed for each bead. **(E)** Raw, unique interaction counts (*y*-axis) of a representative bead with all its interacting neighbors ranked in descending order (*x*-axis). Left plot (blue) shows counts for a representative bead from a SCOPE reaction performed on beads in suspension. Right plot (red) shows counts for a representative bead from a SCOPE reaction performed on beads in a 2D array. **(F)** Mean cumulative distribution function (CDF) of proportion of all interaction counts explained by the top neighbors of a given bead, for SCOPE reactions performed on beads in suspension (blue) or 2D array (red). Gray shadings show the range of CDFs for the 1000 randomly sampled beads from which the mean CDFs were derived. **(G)** Network of interactions of two representative beads (red) from SCOPE reaction performed in 2D array, with and among their top ten neighbors. **(H)** Proportion of read counts from interactions between each bead of interest and their neighbors are overlaid with 100 simulation instances from the inverse square model. **(I)** Region of SCOPE array imaged after staining for two specific “fiducial” sub-barcodes at position 2 (red) or position 4 (blue). Scale bar denotes 100 µm. **(J)** SCOPE reconstruction highlighting the positions of sub-barcodes matching the FISH probes used for imaging. **(K)** Point matching search performed between segmented imaging fiducials and SCOPE reconstruction fiducials. Left: Purple-colored points (beads in SCOPE reconstruction bearing sub-barcode BC2-#33) were scaled, translated, rotated and point-matched onto red-colored centroids from imaging. Right: The same transformation was applied to green-colored points (beads in SCOPE reconstruction bearing sub-barcode BC4-#91) and these were matched to blue-colored centroids from imaging. **(L)** Mean point matching error for both sub-barcodes during gradient descent optimization. Rigid transformations were optimized according to the error for beads with sub-barcode BC2-#33 (red), with sub-barcode BC4-#91 positions (green) held out.

We next sought to evaluate whether SCOPE-derived count matrices are spatially informative. For this, we performed massively parallel sequencing of sender-receiver messenger chimeras recovered from SCOPE reactions performed on either (i) a 2D array of beads; or (ii) beads in suspension. If performing SCOPE results in proximity-dependent reactions between messenger-embedded barcodes derived from physically adjacent beads, then the count matrices of these two reactions should vastly differ. Indeed, for beads in a 2D array, the top 6 partners of a given bead accounted for 36.9% (s.d. 22.5%) of interactions, and the top 20 for 50.1% (s.d. 19.8%). However, for beads in suspension, the top 6 partners of a given bead accounted for only 4.3% (s.d. 12.7%) of interactions, and the top 20 for only 8.9% (s.d. 13.1%) (**Fig. 2E-F**). For the count matrix derived from a 2D array, individual beads and their top interaction partners formed dense subgraphs akin to tightly knit social networks (**Fig. 2G**). Taken together, these results confirm that the count matrices obtained by deep sequencing of sender-receiver messenger chimeras derived from SCOPE reactions performed on 2D arrays are spatially informative.

### SCOPE interaction counts enable spatial reconstruction of fiduciary barcodes

Ranked per-bead interaction count proportions of 2D SCOPE arrays were well-modeled by an inverse square function (**Fig. 2H**). We leveraged this to simulate a diffusive process for molecules derived from beads arrayed in a hexagonal lattice, flexibly parametrized with empirical distributions of the “total sent” and “total received” messengers of any given bead. We then trained a random forest regressor on simulated SCOPE data to convert sparse bead-bead interaction counts into pairwise distances. Training data were generated by simulating bead arrays with known ground-truth positions and modeled interaction profiles, with the simulation re-parameterized for each experimental dataset to match its empirical distributions (**Methods**). Put another way, the regressor is retrained on a dataset-specific simulation before use, to ensure that the count–distance relationship it learns is appropriate for reconstructing the experimental data at hand.

The values in the resulting pairwise distance matrix are inversely related to the counts in the sparse interaction matrix, with “zeros”ーpairs of beads with no detected interactionーassigned a single “long distance” value. Although this approximation violates the triangle inequality at long distances, we posited that the dense local interactions between beads and their closest ring of neighbors would be best resolved with algorithms such as t-SNE or UMAP, which prioritize the preservation of local distances (McInnes et al., 2018; van der Maaten and Hinton, 2008). This was in fact the case, as particularly for smaller arrays, both t-SNE and UMAP accurately reconstructed local communities from a pairwise distance matrix generated by the aforedescribed procedure (**Fig. S3A**).

All rigid transformations (*i.e.* translations, rotations and inversions) are equivalent with respect to the expected distribution of proximity-dependent interactions. As such, rigid transformations present a challenge when comparing the output of a spatial reconstruction to a ground truth image. To evaluate whether fiducial beads could anchor SCOPE reconstructions, we first performed simulations in which an arbitrary sub-barcode (expected to occur in ∼1/96 beads at random) was designated as the fiducial class within simulated bead arrays, and their positions flagged in the ground truth image prior to export. After simulating SCOPE interaction count matrices from these arrays, we applied our reconstruction pipeline to generate inferred bead coordinates. Next, registration of the reconstructed vs. ground truth images was performed by iteratively optimizing the parameters of a rigid transformation using gradient descent. At each iteration, an implementation of the Jonker-Volgenant algorithm for unbalanced assignment was used to match points between the fiducial sets, with the mean Euclidean error between matched pairs serving as the loss function. With sufficient sampling depth (>200 UMIs per bead), reconstructions on a simulated array of 40,000 beads achieved near-perfect alignment to ground truth images, with a mean Euclidean error of 2.19 bead diameters (**Fig. S3B-C**).

Encouraged by these simulations, we sought to integrate fiducial alignment into the SCOPE workflow. For this, we performed fluorescence in situ hybridization (FISH) on a 2D SCOPE array using probes targeting two sub-barcodes (arbitrarily designated red [BC2 #33] and blue [BC4 #91], each expected at random in ∼1/96 beads), followed by imaging (**Fig. 2I**). After running the SCOPE reaction and reconstructing bead coordinates from sequencing-based proximity data, we identified red fiducial barcodes by sequence (**Fig. 2J**). These reconstructed fiducial points were then aligned to their imaged counterparts by optimizing a linear transformation using the same gradient descent point-matching procedure as used on simulations. To assess accuracy, we computed mean Euclidean error on the aligned red fiducials themselves (**Fig. 2K**, left), and separately on the held-out blue fiducials after applying the learned transformation (**Fig. 2K**, right). After alignment, the mean error was 187 pixels (280 µm) on the aligned red points and 179 pixels (269 µm) on the held-out blue points. Over the course of gradient descent, reductions in the mean error for red fiducials was mirrored by similar reductions in the mean error of the held-out blue fiducials (**Fig. 2L**), supporting the conclusion that the alignment procedure is working as intended.

### Computational approach to automated reconstruction of large SCOPE arrays

Although our reconstructions of simulated SCOPE data by “out-of-the-box” application of either UMAP or t-SNE were highly accurate, we encountered various challenges upon either moving to experimental data or upon attempting to scale reconstructions to much larger numbers of beads. To address these challenges and enable reconstruction of large SCOPE arrays from real data, we developed an automated pipeline that integrates data preprocessing, missing data imputation, manifold learning, and parameter optimization.

For data preprocessing, we sought to implement procedures that remove sources of noise specific to SCOPE experiments. First, we remove <u>barcode collision</u>s—instances where a single DNA barcode spuriously appeared in two spatially distinct neighborhoods—using Leiden clustering to identify such nodes as mixed-community outliers (**Fig. S4A**). This approach showed high precision and recall in simulations and removed the expected number of collisions in experimental datasets (**Fig. S4B-E**). Second, we prune <u>spurious</u> <u>edges</u>—strong but biologically implausible interactions linking beads with non-overlapping neighborhoods—using the MinIPath algorithm (Kloosterman et al., 2024). This eliminates “short circuits” that otherwise distort embeddings. Third, we compute the graph’s <u>k-core</u>—the largest subgraph such that all nodes have a minimum degree of k. By setting k to 100, we retain only beads that are well connected and information-rich, to ensure downstream inference is operating on robust subgraphs.

Next, we partition the remaining beads into manageable subsets for downstream analysis. To do this, we apply Leiden clustering to the bead–bead interaction graph, followed by iterative merging of smaller clusters based on shared graph edges. This procedure results in clusters composed of 300 to 2,500 beads. Within each cluster, missing bead-bead interaction values are imputed with a single long-distance value, and all non-zero interactions in the overall pairwise matrix are mapped to distances using our random forest regressor parametrized on simulated data, as described above. This processed pairwise distance matrix is then used to precompute a k-nearest neighbor (k-NN) distance matrix (k = 250). To initialize the approximate spatial relationships among these clusters, we use the PAGA algorithm to generate a force-directed layout of the clusters (39). Finally, the spatial positions of cluster centroids in the PAGA layout, together with 2D-gaussian jitter, serve to initialize a UMAP reconstruction of the full array, with the precomputed k-NN bead-bead distance matrix as input.

Our initial tests showed that hyperparameter selection was important for success. Thus, we implement a grid search over the “min_dist” and “repulsion_strength” UMAP hyperparameters in order to refine the solution. We defined two metrics for hyperparameter selection, <u>contiguity</u> and <u>uniformity</u>. To evaluate contiguity, we generate a binary tiled mask of the point cloud followed by segmentation, and only accept hyperparameter combinations that yield a single connected object. To evaluate uniformity, we rank hyperparameter combinations based on the extent to which they yield a uniform spacing of beads. The hyperparameter selection process is integrated with the aforedescribed workflow, resulting in a fully automated pipeline for spatial reconstruction from SCOPE data.

### Accurate reconstruction of 2D shapes with SCOPE

After developing and validating our reconstruction algorithm based on simulated data, we sought to apply it to experimentally generated datasets. As a first test, we generated a circularly shaped 2D array of SCOPE beads (**Fig. 3A**; 7 mm diameter). As a circle is symmetric, we also introduced fiducials by spotting microliter droplets of poly-dA ssDNA oligos, complementary to the poly-dT cap on decoder oligos (**Fig. 1A**), in a predetermined pattern (12 o’clock, 3 o’clock, 6 o’clock). Following the SCOPE reaction, massively parallel sequencing of chimeras, and application of the aforedescribed computational reconstruction pipeline, we obtained an optics-free reconstruction of the inferred spatial positions of 62,884 beads. Remarkably, although our algorithm does not specify or constrain the shape that the global reconstruction will take, we obtained a circle (**Fig. 3B**). Furthermore, upon mapping the sequenced barcodes corresponding to fiducial-decoder chimeras onto the reconstruction, we obtained a pattern consistent with expectation, *i.e.* diffusion in the vicinity of the 12 o’clock, 3 o’clock and 6 o’clock positions, enabling orientation of the circle (**Fig. S5**).

**Figure 3:**
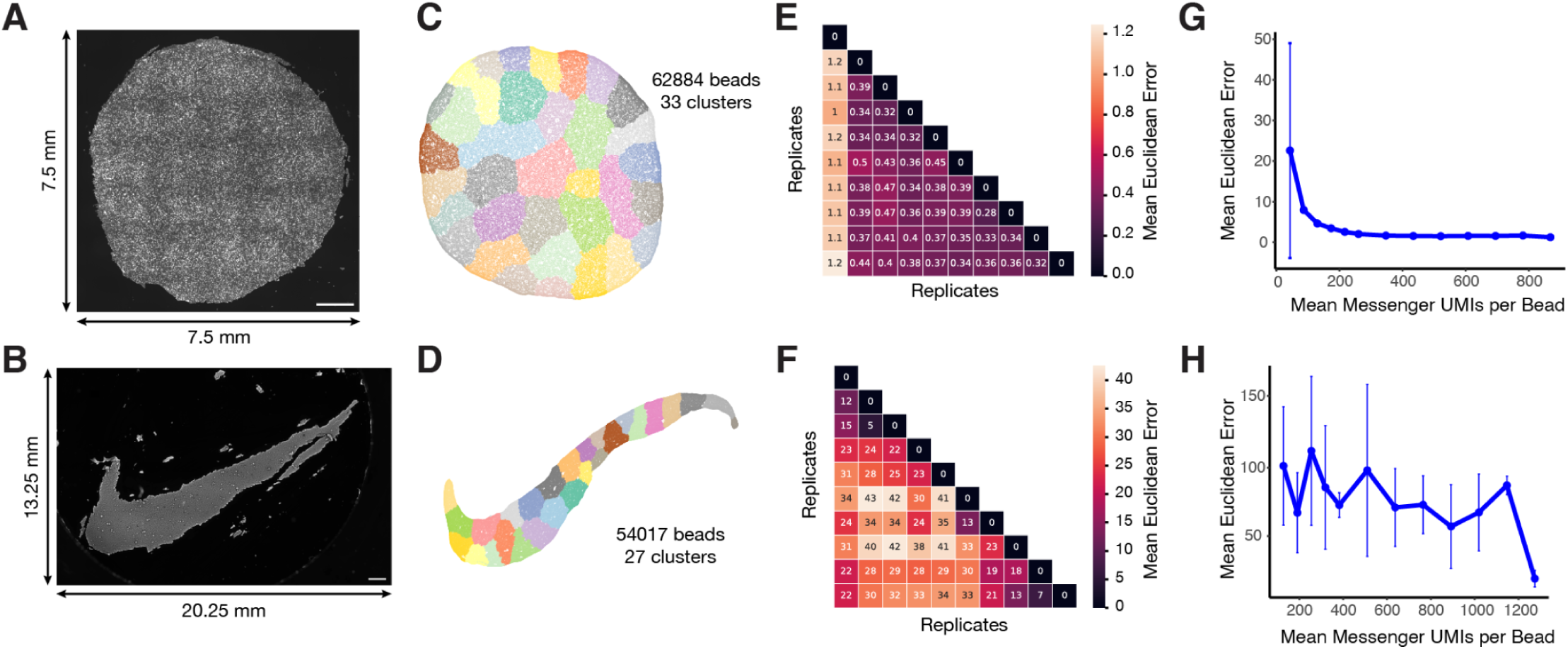
Optics-free reconstruction of 2D shapes with SCOPE. **(A)** Single channel image of a manually cut circular bead array with a 7 mm diameter subjected to hybridization of a sequencing primer followed by single nucleotide extension with dye-conjugated, reversibly terminated dNTPs. Scale bar denotes 1 mm. **(B)** Computational reconstruction of the circular bead array shown in panel **A** with SCOPE. Beads are colored by cluster labels, computed by Leiden clustering on the bead-bead interaction graph. **(C)** Single channel image of a bead array manually cut to an asymmetric “swoosh” shape and stained with SYBR Gold. Scale bar denotes 1 mm. We estimate that the swoosh shape occupies 44 mm^2^. **(D)** Computational reconstruction of the asymmetric swoosh-shaped bead array shown in panel **C** with SCOPE. Beads are colored by cluster labels, computed by Leiden clustering on the bead-bead interaction graph. **(E)** Pairwise mean Euclidean distance between each of 10 independent runs of the reconstruction algorithm on the circular bead array after alignment to a common reference frame. Units are in bead diameters. **(F)** Same as panel **E**, but for asymmetric swoosh-shaped bead array. **(G)** Mean Euclidean bead positioning error for reconstruction of the circular bead array with downsampling of messenger UMIs per bead. Error bars correspond to the range, and points to mean, for 5 reconstructions of each downsampling trial. **(H)** Same as panel **G**, but for asymmetric swoosh-shaped bead array.

Next, we sought to evaluate whether we could apply SCOPE to reconstruct an asymmetric shape. For this, we fabricated an array of SCOPE beads and cut it into the shape of a “swoosh” resembling the Nike logo (16.75 mm x 9.25 mm) (**Fig. 3C**). Following the SCOPE reaction, massively parallel sequencing of chimeras and computational reconstruction, we obtained an optics-free reconstruction of the spatial positions of 54,017 inferred beads that resembles the Nike swoosh (**Fig. 3D**).

Since the Leiden clustering step, which impacts PAGA initialization, is stochastic, and furthermore given that the UMAP algorithm itself is stochastic, our computational reconstruction heuristic gives slightly different results with each run. To evaluate its sensitivity to this, we performed 10 independent computational reconstructions on the data corresponding to both the circular and swoosh arrays (**Fig. S6A,B**). To facilitate alignment and comparison between reconstructions, the median bead-bead lattice edge length was rescaled to 1, and then each independent reconstruction was aligned to the original reconstruction using rigid transformations including rotation and reflections across both axes (**Fig. 3B,E**). After alignment to this common reference frame, we computed the mean Euclidean error between each pair of reconstruction replicates, a metric that averages the positioning error across matched pairs of beads. These results showed that reconstructions of the circular array were highly reproducible (mean error of 0.53; s.d. 0.31 bead diameters), while the reconstructions of the swoosh were more variable (mean error of 28; s.d. 9.2 bead diameters).

Next we investigated the relationship between sequencing depth and reconstruction error. For this, we downsampled the total bead-bead interaction counts in the circular array, retaining 5% to 100% of the total counts. For each downsampling trial, we performed 5 reconstructions and computed the average bead positioning error relative to reconstruction without downsampling (**Fig. 3E,G**). These results indicated that a sequencing depth of ∼400 interactions per bead was sufficient to achieve reconstruction of the symmetric array (mean error of 1.54; s.d. 0.04 bead diameters). However, when the same downsampling rubric was applied to the asymmetric swoosh, error worsened upon even modest downsampling (**Fig. 3F,H**). This suggests that shapes with complex geometries, *e.g.* curves and irregular edges, may require more sequenced interactions per bead for accurate reconstruction. However, the error function used here may be too stringent for such geometries, as modest differences in curvature are exacerbated by a Euclidean error function. For example, a slight error in the curve of the swoosh shape could propagate positioning errors to all beads affected by this partial angular rotation, though it could be argued that geometrically it could all be considered the same error.

### An eye examination for SCOPE reconstruction of 2D images

The resolving power of optical systems is constrained by the wavelength of light and the numerical aperture of their optics. In contrast, DNA-based spatial reconstruction should be limited by diffusion kinetics rather than optical physics. Although these experiments remain well above the resolution limits of microscopy, we next sought to test whether SCOPE could reconstruct large images while preserving scale across multiple resolutions. We also wanted to determine whether it could resolve distinct features—effectively moving from bead-defined shapes to decoder-defined “color” images. To this end, we performed a virtual eye examination, using SCOPE to reconstruct elements of the classic Snellen eye chart for visual acuity.

Using a microarray printer, we deposited picoliter droplets of 13 poly(A)-tailed oligonucleotide “paints” (100 µm spot size; 50 µm center-to-center pitch) onto a rectangular 2D SCOPE array measuring 17.18 mm × 40.97 mm (704 mm^2^). Three paints were fluorescently labeled, enabling orthogonal visualization by microscopy (**Fig. 4A**), which revealed fine striations corresponding to the printer’s line stepping (**Fig. 4A**, insets). Imaging-based estimates indicated that each printed oligonucleotide spot covered an average of ∼45 beads (s.d. = 1).

**Figure 4:**
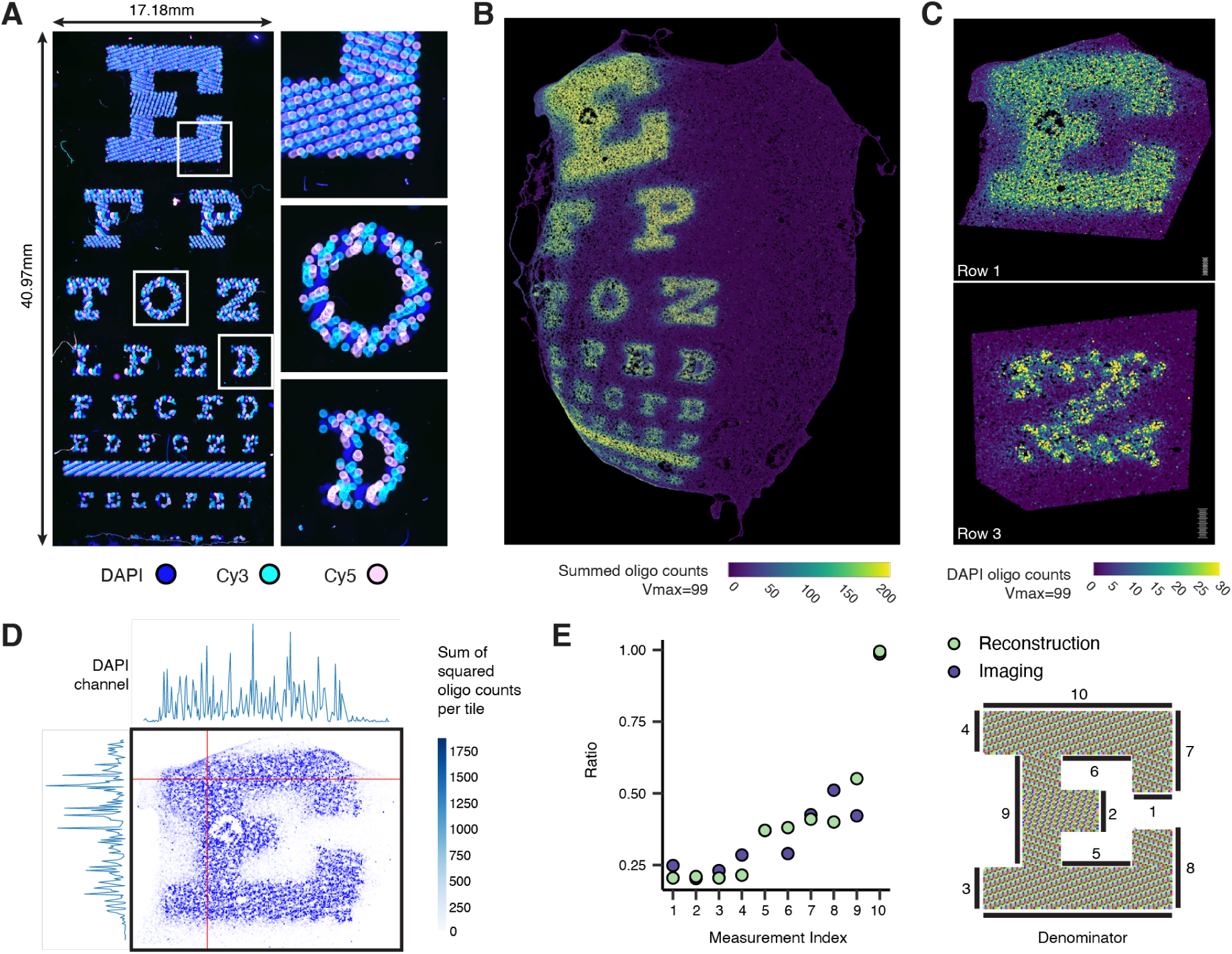
An eye examination for SCOPE. **(A)** Composite fluorescent image of a Snellen eye chart for visual acuity printed onto a 2D SCOPE array using a microarray printer. Of the 13 printed oligonucleotides (“paints”), three were fluorescently labeled—either via dye conjugation (Cy3, Cy5) or inclusion of DAPI in solution—allowing direct microscopic visualization. Insets show the “E” (top), “O” (middle), and “D” (bottom) for reference. **(B)** Maximum-intensity projection of the reconstructed array comprising 1.44 million beads, colored by summed decoder counts across all paints. **(C)** Maximum intensity projection of a single oligo channel (DAPI channel oligo) from the “E” in Row 1 and the “Z” in Row 3. The region corresponding to each letter was cropped from the full reconstruction, rotated, and displayed at different scales with a common ruler for reference. **(D)** Locally binned image showing the sum-of-squared oligo counts per bin, with horizontal (top) and vertical (left) line intensity profiles corresponding to the red crosshairs. (**E**) Dimensional fidelity of the reconstructed “E.” Plotted points show the ratio between each of 10 feature lengths indicated by bars at the right and the length of the base of the E, measured via optics (“imaging”) or SCOPE (“reconstruction”) using ImageJ.

After the SCOPE reaction, we generated two sequencing libraries: (i) messenger chimeras encoding bead-bead proximities, and (ii) decoder chimeras mapping the positions of the microarray-spotted paints. Sequencing of messenger chimeras identified DNA barcodes corresponding to 1.44 million beads and 2.55 billion unique bead-bead interactions. Sequencing of the accompanying decoder library linked an average of 52 (s.d. = 65) poly(dA)-tailed “painted” molecules to each bead. Applying the computational pipeline described above, we mapped the positions of paints onto the reconstructed 2D array. However, the initial 2D UMAP reconstruction exhibited macroscopic distortions, potentially caused by non-uniform diffusion across this large array during the SCOPE reaction (**Fig. S7**). To mitigate such effects, we adopted a two-stage embedding approach: first computing a 3D UMAP from the precomputed k-nearest neighbor distance graph, then “flattening” the manifold to 2D by generating a second UMAP that used the 3D coordinates as input. This approach yielded a global map consistent with a reaction confined to a planar slide. We then summed decoder counts for each paint at each position and visualized the resulting image (**Fig. 4B**).

With this revised heuristic, the SCOPE reconstruction reproduced the Snellen chart with substantially reduced macroscopic distortion (compare **Fig. S7** vs. **Fig. 4B**). However, it remained offset to the left, which may be due to off-center printing during microarray deposition. Residual warping, which gives the appearance of a weakly curved surface, potentially reflects diffusion dynamics near the edges of the bead array—specifically, a radial outward bias analogous to capillary flow in an unconfined film. Future refinements, whether experimental (*e.g.* imposing a physical barrier to mitigate radial outward bias), computational (*e.g.* modeling and subtracting such biases), or both, may help to further improve the fidelity of large image reconstruction.

The Snellen chart assesses visual acuity through letters arranged from largest to smallest. Distinct striated patterns—mirroring those observed by microscopy (**Fig. 4A**)—were recovered through at least the third row of letters when each oligonucleotide “paint” channel was visualized individually, *e.g.* within the “E” of the first row (**Fig. 4C**, top; further confirmed by horizontal and vertical line intensity profiles shown in **Fig. 4D**) and the “Z” of the third row (**Fig. 4C**, bottom).

To further assess fine-scale reconstruction accuracy, we focused on the large “E,” which encompassed 190,790 beads. During reconstruction, SCOPE operates solely on messenger interactions, without access to decoder information. To compare dimensional fidelity between the printed and reconstructed images, we measured the lengths of multiple features within the serif-styled letter E that we had printed, normalizing each feature to the letter’s baseline length. Feature measurements were highly correlated between the optical image and SCOPE-based reconstruction (Pearson’s *r* = 0.96; Spearman’s ⍴ *=* 0.90) (**Fig. 4E**), indicating that SCOPE can recover both local and global geometric scales in the micron to centimeter range while preserving proportional relationships across the image.

### Extending SCOPE to the optics-free reconstruction to 3D volumes

Inspired by the observation that a three-dimensional UMAP embedding improved reconstruction accuracy for large 2D arrays, we next asked whether SCOPE could reconstruct truly three-dimensional structures. To test this, we embedded 100 µm DNA-barcoded beads within an acrylamide hydrogel matrix by polymerizing the gel via a dense bead slurry cast into silicone molds of predefined shapes. Specifically, we used 3D molds of a teddy bear, star, butterfly, and block letter (“E”), each with volumes on the order of 75-100 mm^3^. Following polymerization, SCOPE proximity reactions captured diffusion-limited hybridization events between barcodes on spatially adjacent beads, generating chimeric molecules that encode three-dimensional neighborhood relationships. These were sequenced and processed analogously to the 2D bead array experiments to yield bead–bead interaction matrices. Applying a 3D implementation of UMAP, coupled with a grid search over key parameters, we inferred bead positions from these interaction data. Encouragingly, the resulting embeddings recapitulated the overall geometry of each of the four original molds (**Fig. 5A–D**).

**Figure 5:**
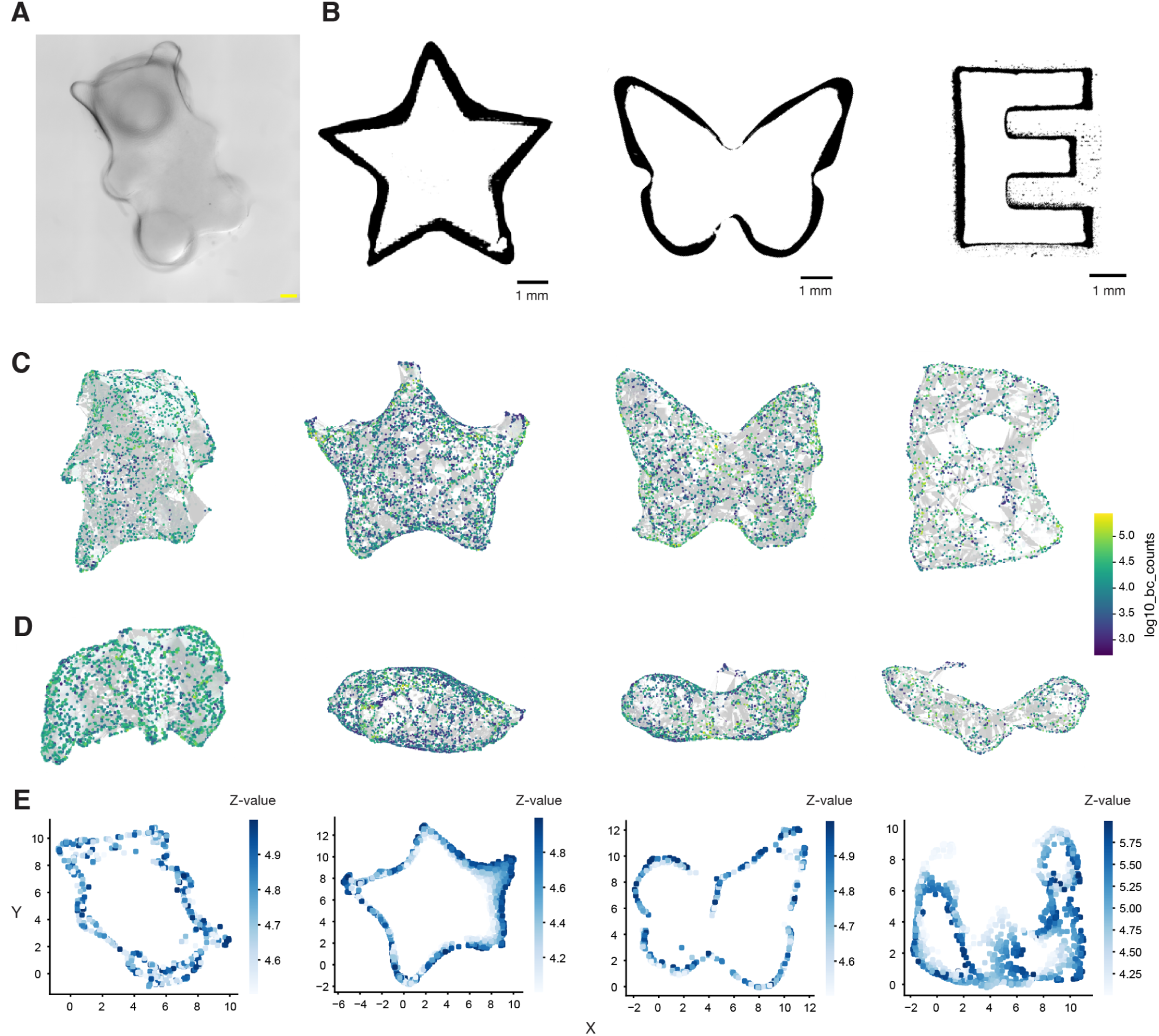
UMAP reconstructions of 3D volumes with SCOPE. **(A)** Brightfield image of a bead-embedded polyacrylamide gel matrix from a teddy bear mold. Yellow scale bar denotes 1 mm. **(B)** Binary masks of brightfield images of the silicone molds used to create bead-embedded 3D gels for star, butterfly and letter E-shaped molds. The depth of each mold is 2 mm. Black scale bars denote 1 mm. **(C)** Top-views of 3D UMAP reconstructions of the bead-embedded gels from the bear, star, butterfly, and letter E-shaped molds, using n_neighbors=15, min_dist=0.4, and 1,000 training epochs on the interaction matrix. Each barcode was normalized to 10,000 total counts per barcode and then log-transformed. Points are bead barcodes in 3D space overlaid on a concave hull (white) calculated by the Python package “alphashape” and colored by the log10 UMI count of each barcode. Reconstructions have been rotated to show the front view. **(D)** Same as panel **C**, but side-views. **(E)** Z-axis cross-sections of 3D UMAP embeddings from bead-embedded hydrogel reconstructions. Each slice shows reconstructed bead positions at successive depths in the inferred volume. The absence of points in central regions highlights the hollow character of computational reconstructions, consistent with reduced recovery of interior beads due to diffusion or steric limitations during SCOPE reaction.

Exploration of UMAP parameter dependencies informed the reconstruction process (**Figs. S8, S9**). Increasing the number of training epochs improved convergence and shape fidelity, while moderate *min-dist* values (≈0.4) preserved local continuity without excessive fragmentation or inward curling. The “cosine” distance metric yielded more stable embeddings and eliminated the need to densify the sparse interaction matrix. Normalization strategies applied prior to manifold learning had smaller effects on reconstruction quality; we ultimately used a count matrix normalized to 10,000 total counts per barcode and then log-transformed. Collectively, these observations underscore the importance of careful parameter tuning for achieving accurate and stable 3D reconstructions.

Although the 3D embeddings recovered by SCOPE captured the global outlines of each molded shape, cross-sectional visualization revealed that their interiors were largely hollow (**Fig. 5E**), indicating underrepresentation of beads from the interior of each mold. We attribute this to a recovery limitation rather than a shortcoming of the reconstruction algorithm. Unlike volumetric DNA microscopy, in which the hydrogel-embedded specimen is fully digested to liberate molecules throughout the volume for downstream sequencing (Qian et al., 2026), the current implementation of SCOPE leaves the polyacrylamide hydrogel intact. As a result, both the formation and recovery of chimeric molecules are diffusion-limited: enzymes must access the interior of the scaffold to cleave bead barcodes, cleaved barcodes must diffuse locally to encounter neighboring beads, and the resulting chimeric products must diffuse out of the scaffold to be recovered for sequencing. Each of these steps likely biases against molecules generated in the interior of the sample. To isolate the diffusion problem, we performed a control experiment in which barcoded beads were allowed to settle in solution at the bottom of a tube without a polymerized hydrogel scaffold, and SCOPE proximity reactions were performed directly on the settled beads. 3D UMAP reconstruction of these data yielded a solid, non-hollow structure consistent with the expected conical geometry of a tube bottom (**Fig. S8A-B**), indicating that SCOPE can recover volumetric structure when recovery is not diffusion-limited.

Because of these missing interior regions, the quantitative metrics developed for evaluating 2D reconstructions were not directly applicable to the 3D embeddings. Lowering the UMI inclusion threshold from 1,000 to 100 slightly increased interior density but introduced low-confidence bead clusters inconsistent with the known geometry (**Fig. S9**), and so did not provide a usable rescue. We therefore selected final embeddings from the range of hyperparameter combinations explored (**Figs. S10, S11**) based on subjective resemblance to the expected global geometry of each molded shape.

Although further work will be required to address these limitations, these experiments establish that SCOPE can recover the global geometry of three-dimensional objects directly from molecular proximity data, laying the groundwork for future volumetric, optics-free spatial reconstructions.

## DISCUSSION

Molecular self-assembly is a defining feature of living systems (Evans et al., 2024). Biological organization arises from simple local interactions among molecules and cells—mostly diffusion-limited—that collectively scale from nanometers to meters, *e.g.* as when a micron-scale zygotic nucleus encodes the blueprint for a 30-meter blue whale. The same principle underlies DNA microscopy and related approaches, which exploit proximity-dependent reactions to infer spatial relationships without optics. Inspired by these concepts and by the physics of molecular diffusion, we developed SCOPE, which propagates information obtained solely at the micron scale to derive optics-free reconstructions at the centimeter scale—a span of four orders of magnitude.

SCOPE relies on barcoded hydrogel beads that transmit information about their identity to neighboring beads through highly localized, diffusion-limited DNA-DNA proximity reactions. We show that this form of pairwise interaction data can be computationally transformed to reconstruct shapes, images, and volumes of arbitrary size, given sufficient sampling. A key feature is that SCOPE reactions occur in constant experimental time regardless of array size. We envision that such self-registering arrays could be broadly useful in spatial genomics, where current state-of-the-art methods still rely heavily on microscopy—either to map barcode sequences to spatial positions or to register molecular signals within intact samples. This dependence on optics imposes an inherent trade-off between physical area and spatial resolution, such that the collection of large-scale datasets (*e.g.* spatial transcriptomics of the whole mouse brain (Langlieb et al., 2023; Zhang et al., 2023)) are major efforts rather than routine experiments.

Independently developed, concurrently reported methods from the Chen (‘imaging-free spatial genomics’) and Cao (‘IRISeq’) groups also infer spatial relationships through diffusion-mediated barcode exchange but differ in architecture and computational formulation (Abdulraouf et al., 2024; Hu et al., 2025). In their approaches, two bead types—“transmitters” and “receivers”—are co-deposited such that oligos from transmitters diffuse and are captured by sparsely distributed receivers. Each receiver’s unique mixture of captured barcodes defines its neighborhood. In contrast, all SCOPE beads act as both transmitters (*i.e.* senders, in our terminology) and receivers, producing a square, asymmetric bead-bead interaction matrix. Computationally, while all three methods employ UMAP to infer bead positions, SCOPE differs in how this manifold learning step is applied. The methods developed by the Chen and Cao groups treat the receiver-sender pairwise interaction matrix as a high-dimensional feature matrix and apply UMAP directly using preselected hyperparameters. In SCOPE, the interaction matrix is instead treated as a noisy measurement of the underlying pairwise distance matrix among beads on the planar slide surface. This matrix is first transformed through simulation and a learnable mapping function to approximate true distances, and UMAP is then applied directly to this distance matrix, with hyperparameters optimized dynamically via grid search. In this way, SCOPE’s use of UMAP arguably aligns more closely with the original design intent of the algorithm: reconstructing the positions of uniformly distributed points on a two-dimensional manifold, without dimensionality reduction.

Although we focused here on reconstructing shapes, images, and volumes where “ground truth” was known, extending SCOPE to capture and decode the relative spatial positions of biomolecules or cells within biologically intact samples is a logical next step. The related work from Chen and Cao (Abdulraouf et al., 2024; Hu et al., 2025), as well as the original DNA microscopy method and its extensions (Qian and Weinstein, 2025; Weinstein et al., 2019), suggests that such biological applications are feasible. For most spatial genomics applications, the typical geometry will likely be a simple rectangular monolayer onto which a tissue section of arbitrary size and shape can be overlaid. However, because SCOPE does not depend on patterned deposition or rigid substrates, it could also be adapted for non-planar geometries such as curved surfaces or three-dimensional scaffolds. Beyond reconstructing 3D bead-embedded gels, bead-laden hydrogel matrices could serve as internal scaffolds that conform to hollow tissues. For example, embedding a bead-packed gel within a tissue lumen—such as the gut—could allow surface beads to capture RNA transcripts from adjacent cells while bead-bead associations define the surface topology. Such an approach could enable simultaneous inference of 3D tissue architecture and spatially resolved transcriptomic profiles.

Experimentally and computationally, what limits SCOPE? Barcoded beads can be produced at exponential scales with split-pool strategies and easily cast to generate large 2D arrays. As noted above, the SCOPE reaction occurs in constant experimental time regardless of array size, and even for very large arrays, reaction volumes remain trivial. At smaller scales, stochastic variation in local diffusion or hybridization efficiencies—and potential anisotropy in diffusion through heterogeneous media—may introduce noise in the inferred local distances. In the current implementation, we used 20 μm beads as a tractable starting point for developing the method. However, although the effective spatial resolution is not solely limited by bead size alone, it could in principle be improved through a combination of smaller-diameter beads, higher packing densities, deeper sequencing of molecular interactions, optimization of SCOPE reaction conditions and further algorithmic improvements.

For large-scale reconstructions such as the Snellen eye chart, sequencing depth of the chimeric sender–receiver libraries can become a limiting factor. Additionally, for large planar arrays, radial distortions in diffusion rates present another hurdle for faithful macro-scale reconstruction, which could potentially be mitigated through experimental refinements. Finally, in 3D volumetric reconstructions, we also observed gross underrepresentation of interior beads, resulting in reconstructions with hollow centers—likely due to reduced accessibility of inner beads during enzymatic cleavage, ligation, or chimera recovery. Algorithmically, SCOPE reconstructions rely on fast graph-based and manifold-learning methods that scale efficiently to millions of beads, but pushing toward billions will require distributed implementations of the distance mapping and embedding steps to maintain tractable runtimes.

Modern molecular and cellular biology increasingly merges DNA barcoding (*e.g.* UMIs, MPRAs, combinatorial indexing) with spatial anchoring (*e.g.* massively parallel sequencing) to parallelize experimentation within single reaction volumes. SCOPE integrates these principles to reconstruct 2D and 3D spatial information in an optics-free manner. Looking forward, deeper intersections of DNA barcoding and spatial anchoring—both *in vitro* and eventually *in vivo*—may enable new classes of molecular experiments: from molecular computing and massively multiplexed encoding/decoding, to information exchange among cells during development, or even spatially coherent DNA automata.

## Acknowledgments

We thank members of the Shendure and Srivatsan Labs for helpful discussions and reagents. We also thank the Lampe Lab for the use of their array printer, as well as the imaging facilities and high-performance computing facilities at both Fred Hutchinson Cancer Research Center and the University of Washington. Additionally, we thank the Biology Imaging Facility at the University of Washington and the Cellular Imaging Shared Resource, RRID:SCR_022609, of the Fred Hutch/University of Washington/Seattle Children’s Cancer Consortium (P30 CA015704). We disclose that language editing, proofreading and coding were supported by AI-based tools; these were not used for conceptual development or primary manuscript writing. The authors take full responsibility for the contents of this manuscript.

## Funding

This work was supported by the Brotman Baty Institute for Precision Medicine and grants from the Paul G. Allen Frontiers Group (Allen Discovery Center for Cell Lineage Tracing to J.S. and C.T.), Alex’s Lemonade Stand Foundation (Grant 19-15730), Crazy 8 Initiative (to J.S.), and National Institutes of Health (HG010632 to J.S. and C.T.; AI146028 to F.A.M.). H.L. is supported by the NSF Graduate Research Fellowship Program (DGE-2140004). J.S. and F.A.M. are Investigators of the Howard Hughes Medical Institute.

## Author Contributions

S.S., J.S., C.T., and J.G. were involved in the initial planning and conceptualization of the project. S.S., H.L., J.G., M.P.R., and R.M.D. performed the experiments. H.L., Y.H., and O.W. designed and streamlined bioinformatic pipelines. F.A.M. designed and implemented the simulation model. S.K. designed and implemented the reconstruction algorithm with substantial feedback from S.S. Y.H. designed and implemented the doublet detection method. M.C. and S.S. performed segmentation and image analysis. S.S., J.S., H.L., and S.K. wrote the manuscript with input from all co-authors.

## Competing interests

The Fred Hutchinson Cancer Center and University of Washington have filed a patent application partially based on this work, on which S.S., J.S., H.L., J.G. and S.K. are listed as inventors. J.S. is on the scientific advisory board, a consultant, and/or a co-founder of Guardant Health, Phase Genomics, Adaptive Biotechnologies, Sixth Street Capital, Pacific Biosciences, Cellular Intelligence and 10x Genomics. All other authors declare no competing interests. The other authors have no competing interests to declare.

## Data and materials availability

Downsampled sequencing data for the reconstruction experiments have been deposited to Github (https://github.com/SrivatsanLab/SCOPE), while the raw sequencing data and processed data can be accessed via the NCBI Gene Expression Omnibus (accession: GSE313591). Our custom reconstruction pipeline used to reconstruct the SCOPE experiments is available at https://github.com/SrivatsanLab/SCOPE and simulation code https://github.com/matsengrp/sci-space-v2.

## Supplementary Figures

**Figure S1:**
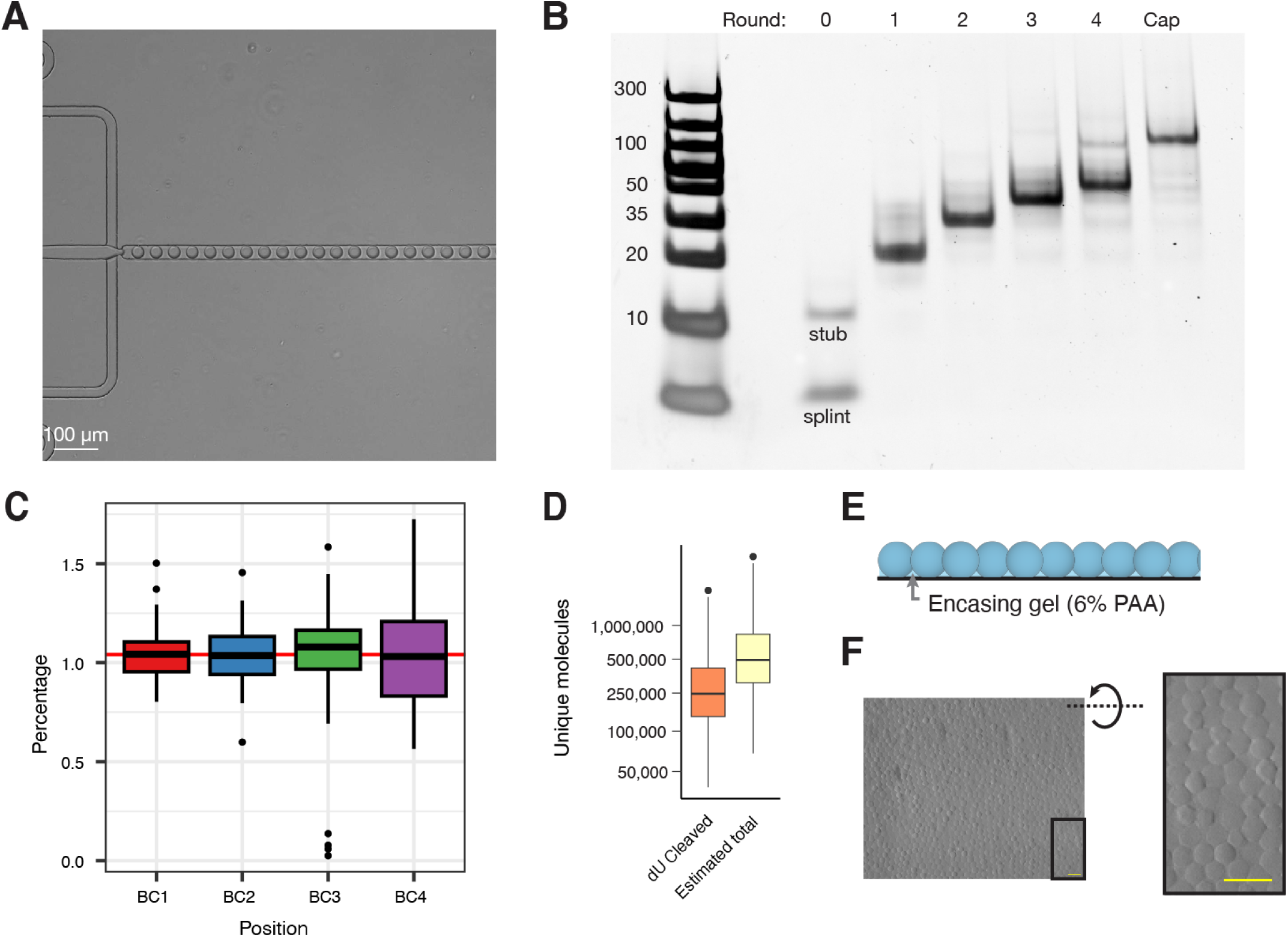
Barcoded hydrogel bead generation and bead array fabrication. **(A)** Image of the flow focusing microfluidics generator while producing polyacrylamide hydrogel beads. **(B)** 10% polyacrylamide gel of oligos released by USER-mediated cleavage from the initial bead (round 0) vs. after each round of DNA synthesis by ligation of the barcode (rounds 1, 2, 3, 4) or functional cap (Cap). Left: DNA size ladder (bp). In the lane corresponding to the initial bead (round 0), the band labeled “stub” corresponds to what is cleaved off the strand that was initially incorporated into the hydrogel bead, while the the band labeled “splint” is a primer that anneals to the stub to create a 4-bp overhang handle for the ligation of the first of four splint barcodes (Delley and Abate, 2021). **(C)** 96 sub-barcodes were used in each of four rounds of the split-pool procedure for barcode generation (Delley and Abate, 2021). Shown in the box plot is the percentage of each sub-barcode incorporated to full barcode at each position. The red line at 1.04% corresponds to uniform incorporation (1/96). **(D)** Based on amplification and sequencing of UMIs associated with barcodes enzymatically released from individual beads, together with the fact that only ∼50% of the barcodes have a dU at their stem and are expected to be cleaved, we estimate that each bead bears 491,811 (interquartile range: 308,757-833,744) functionalized, barcoded oligos. **(E)** Schematic of a monolayer of hydrogel beads inside a 6% (v/v) PAA (polyacrylamide) encasing gel. **(F)** Differential interference contrast microscopy image of beads in an encasing gel. Yellow scale bar: 100 µm.

**Figure S2:**
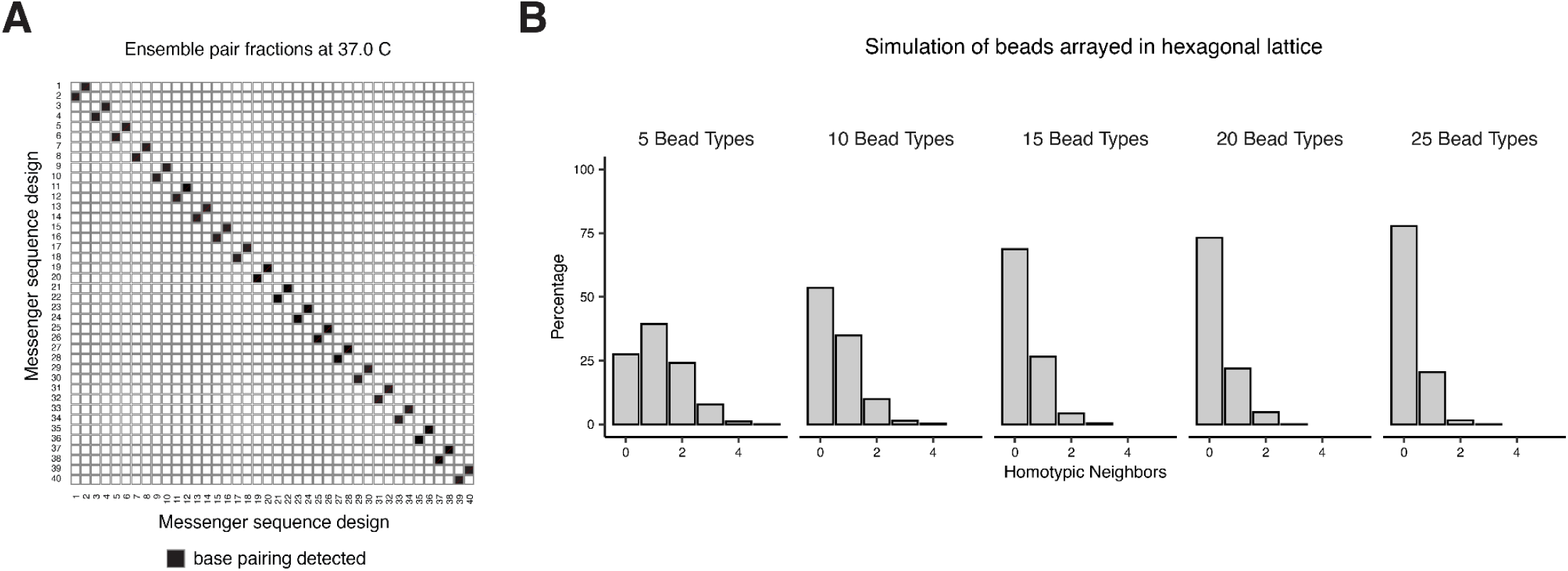
Complementarity and predicted outcomes of using twenty pairs of messenger sequence designs. **(A)** Fractions of oligos predicted to be bound at equilibrium in a pool of 40 designed messenger cap sequences (20 messenger caps x 2 orientations) at 37°C as estimated by Nupack (Fornace et al., 2022). Complementary strands are adjacent in numbering (*e.g.* 1/2, 3/4 … 39/40). **(B)** We simulated the random assignment of a given number of bead subtypes to beads arrayed in a hexagonal lattice and computed, for any given bead, the proportion of immediately neighboring beads that are of the same subtype (*i.e.* homotypic neighbors).

**Figure S3:**
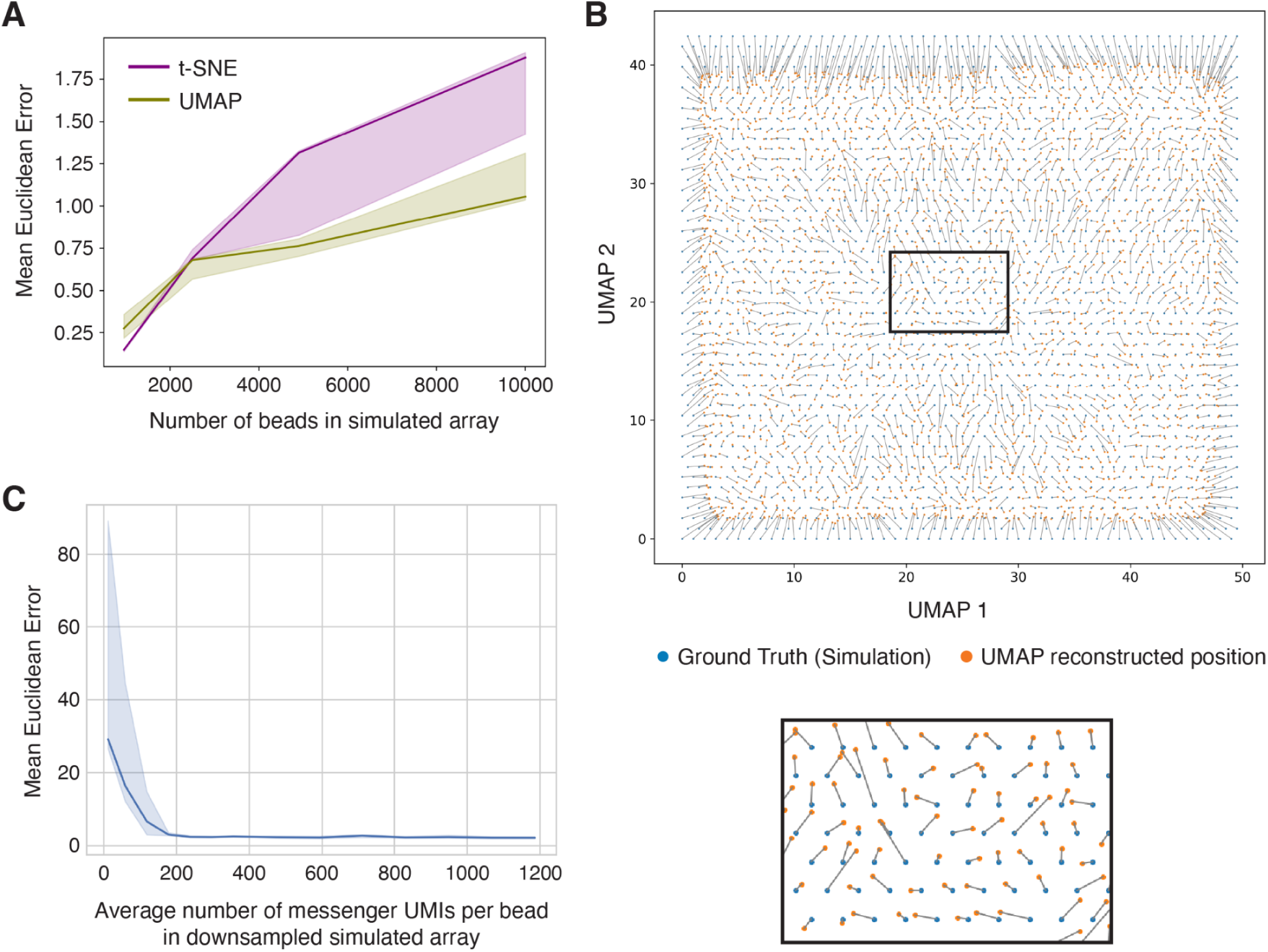
Reconstructing simulated SCOPE bead arrays. **(A)** Mean Euclidean error of UMAP (green) or t-SNE (purple) applied to simulated SCOPE bead arrays of different sizes. Mean Euclidean error is measured as the average difference in distance between an inferred position and the ground truth simulated position after a linear point cloud registration and point matching. The line shows the median of 5 trials and the shaded areas show the interquartile range. Units are in bead diameters. **(B)** A representative simulated rectangular array of 2500 beads reconstructed with UMAP. Lines indicate matching between ground truth and UMAP inferred positions after linear point cloud registration. Bead positioning errors are greater at the boundary of the shape. **(C)** Mean Euclidean error of a simulated and reconstructed 40,000 bead array as a function of simulated sequencing depth. Each point is the median of 10 trials. The shaded blue area shows the interquartile range for the set of trials.

**Figure S4:**
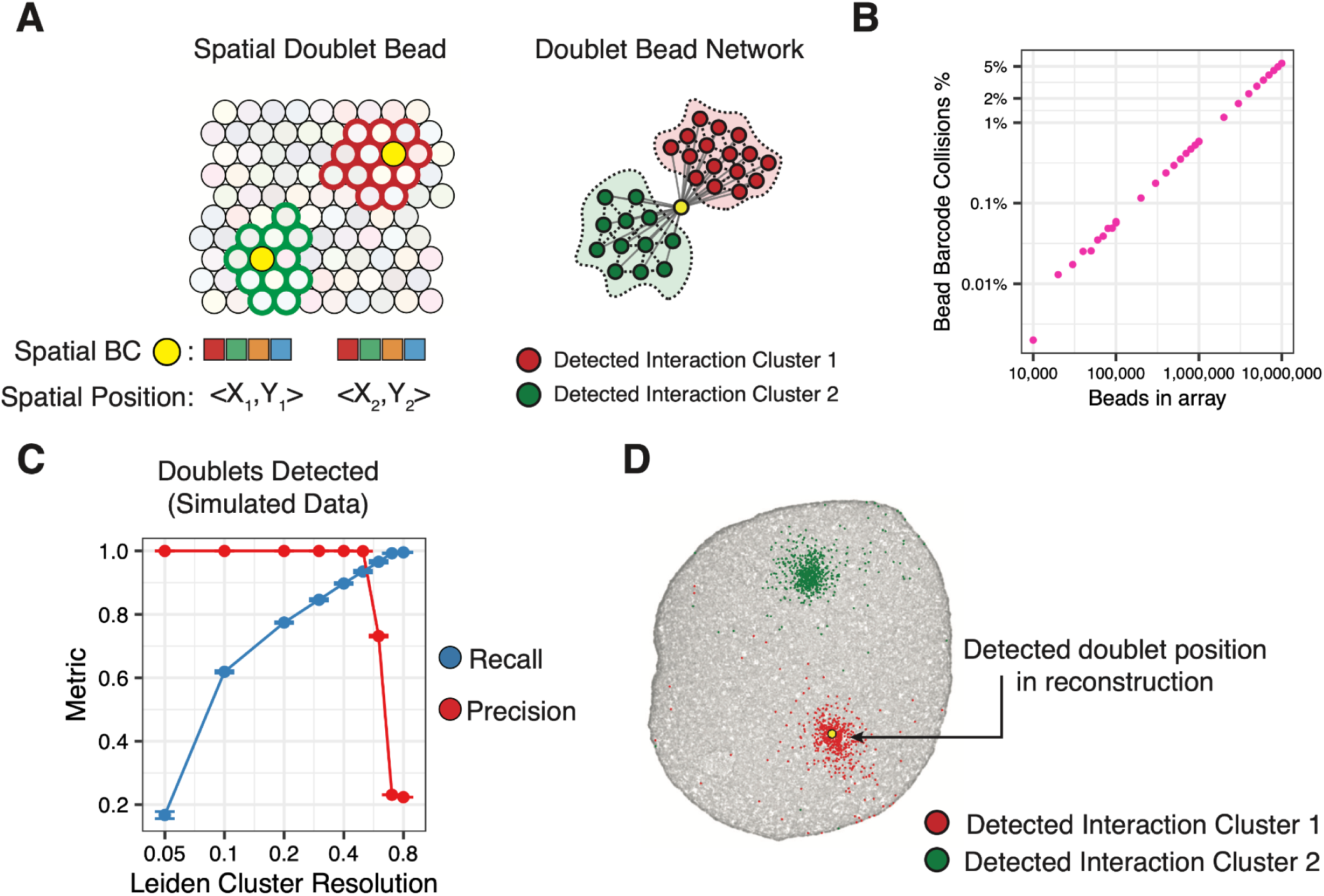
Doublet detection by community detection. **(A)** Doublets are defined as beads that have two spatial positions but share an identical DNA barcode (ground truth – left). In sequencing data these doublets appear as a single bead with collapsed interaction networks. **(B)** Expected rates of doublet occurrence for arrays of a given size. Estimated by computing the birthday problem for a range of array sizes with a fixed set of 96^4^ barcodes. **(C)** Doublets detected in simulation using Leiden clustering and a range of cluster resolutions. Recall and precision are shown with error bars indicating standard deviation. **(D)** Examples of detected doublets during reconstruction of the circular array. Highlighted in red and green are the two communities detected for a single bead (shown in yellow).

**Figure S5:**
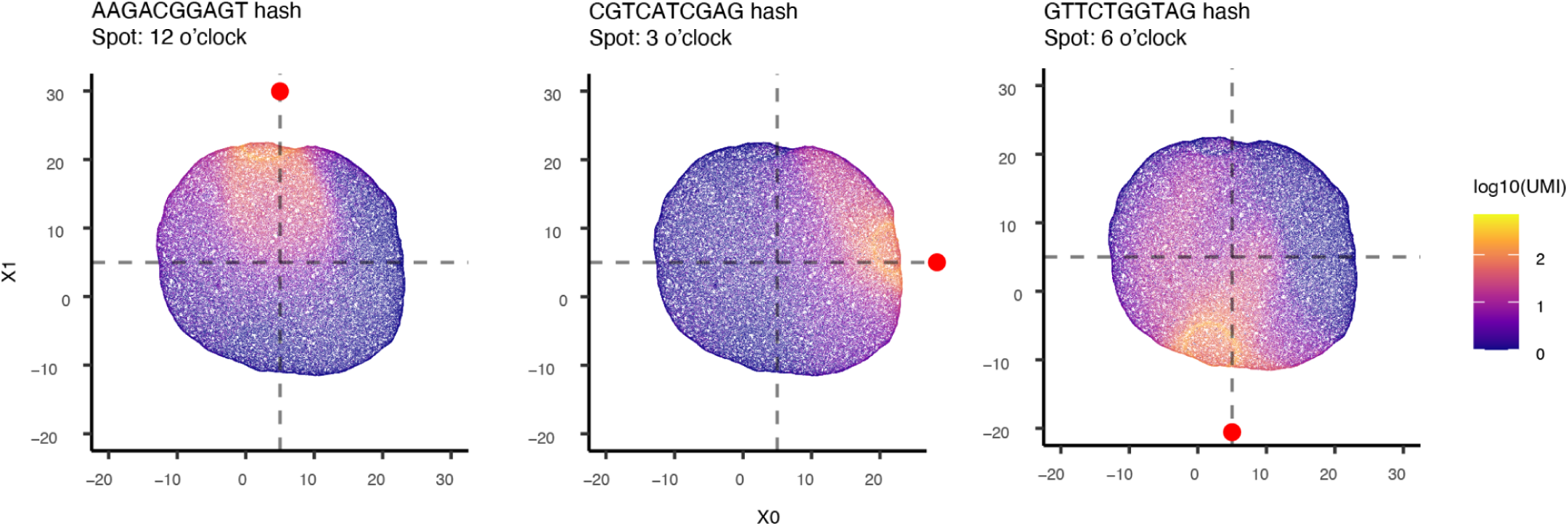
Spotted poly-dA fiducials enable orientation of SCOPE reconstructions of symmetric 2D arrays. Poly-dA–tailed hash oligonucleotides were manually spotted onto the circular bead array at the 12 o’clock, 3 o’clock, and 6 o’clock positions prior to the SCOPE reaction. During decoding, complementary poly-dT–capped decoder molecules captured these fiducials, producing locally concentrated clusters of corresponding barcoded sequences. In the reconstructed array (three versions of which are shown here without rotation or reflection relative to one another), the three fiducial barcodes are locally concentrated at expected relative positions (red dots), confirming correct geometry and providing reference points for orienting symmetric reconstructions. X0 and X1 denote the spatial axes outputted by the reconstruction pipeline.

**Figure S6:**
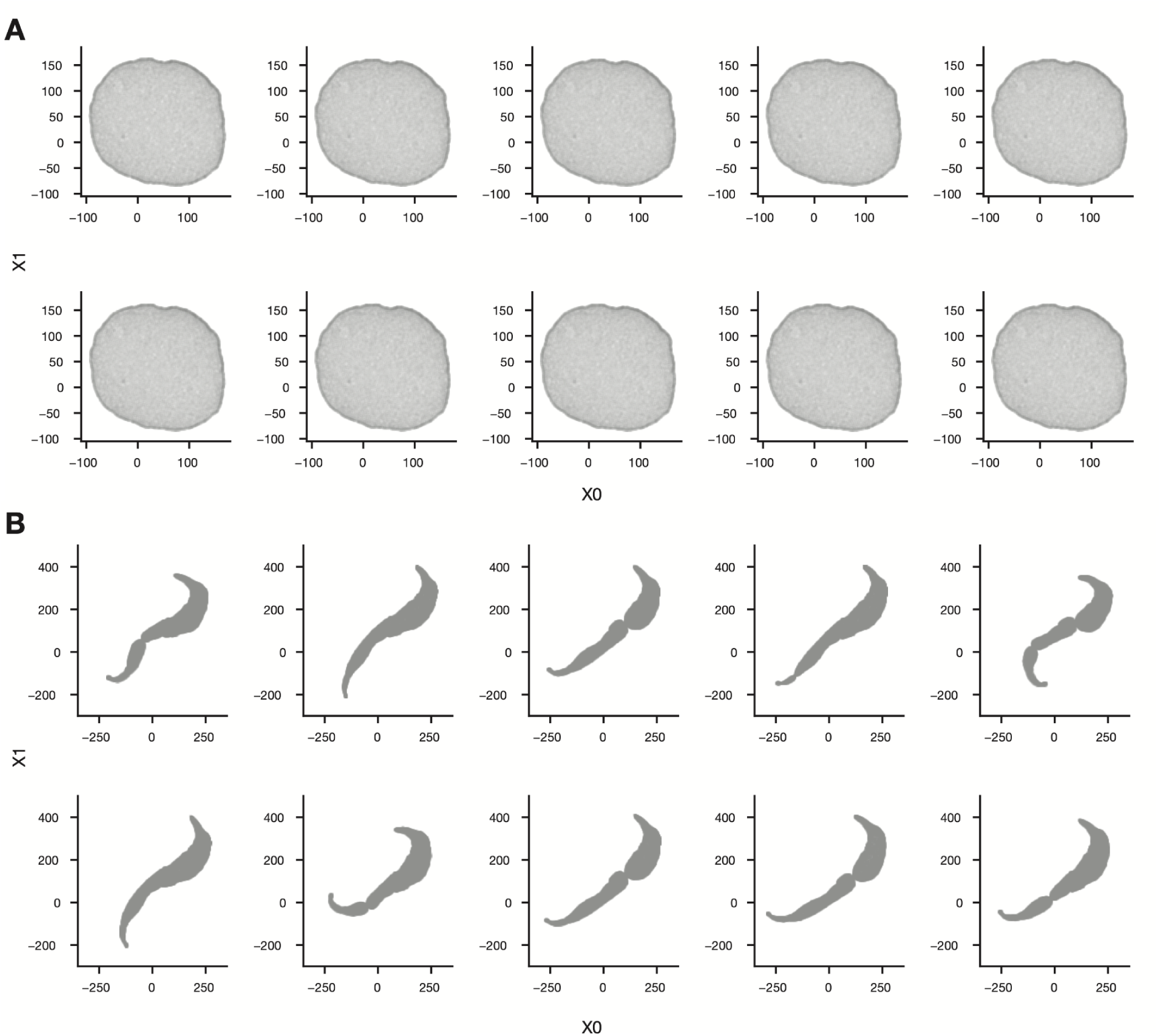
Ten iterations of computational reconstruction of circular and swoosh arrays for **(A)** circular bead array or **(B)** asymmetric “swoosh” array. The pipeline was re-run independently for each reconstruction with identical inputs and initial hyperparameters each time, only varying the random seed. Each reconstruction has been transformed (including rotation, reflection, scaling) for visualization. X0 and X1 denote the spatial axes outputted by the reconstruction pipeline.

**Figure S7:**
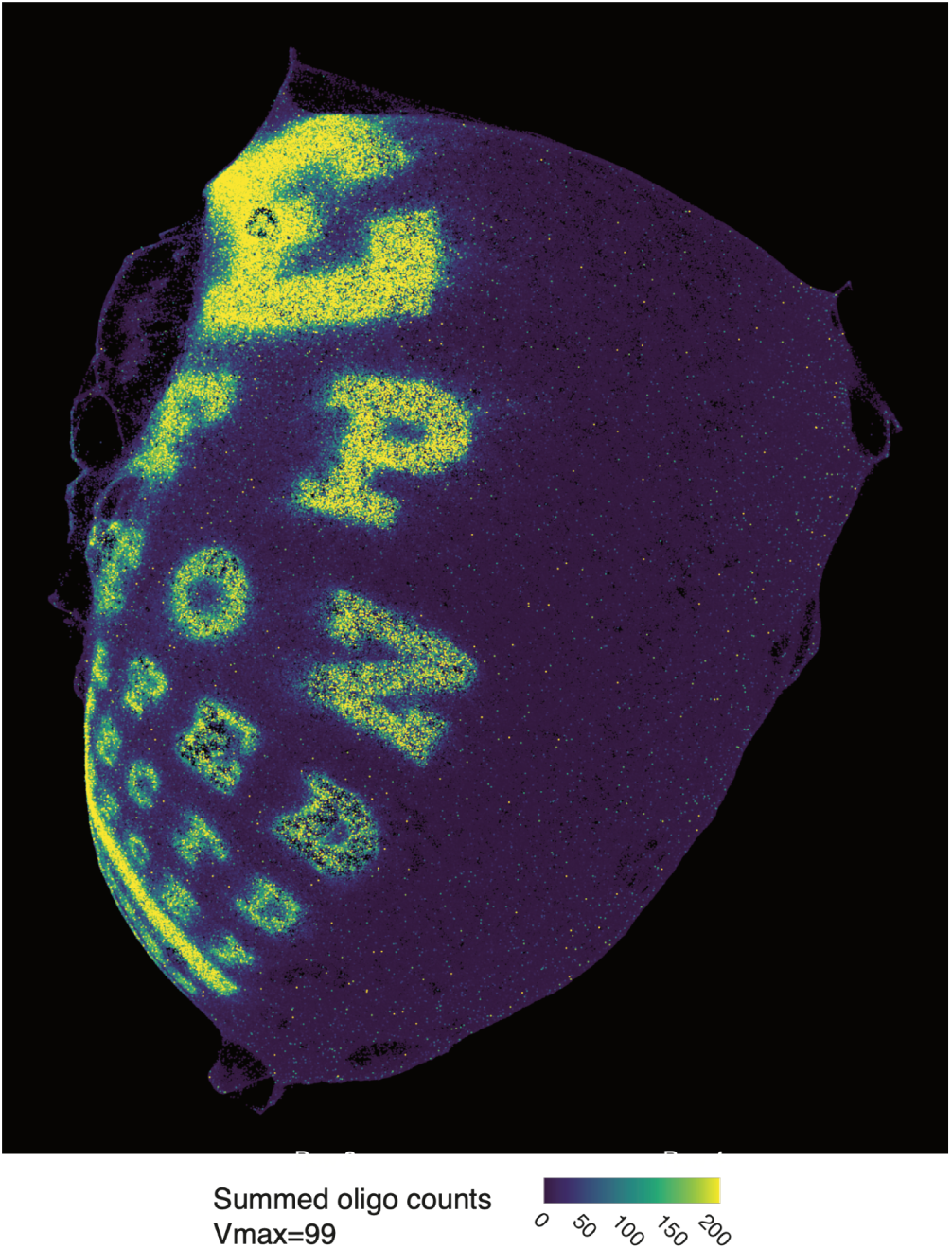
Initial 2D UMAP result for Snellen eye chart experiment prior to 3D flattening. Maximum-intensity projection of the reconstructed Snellen eye chart generated using the standard 2D UMAP pipeline applied directly to the bead–bead interaction matrix. Each point represents a DNA-barcoded bead (n = 1.44 million), colored by the summed decoder counts across all 13 printed poly(A)-tailed oligonucleotide “paints.” Although major features of the eye chart are discernible, the reconstruction exhibits macroscopic warping and curvature, likely reflecting non-uniform diffusion during the SCOPE reaction across this large array.

**Figure S8:**
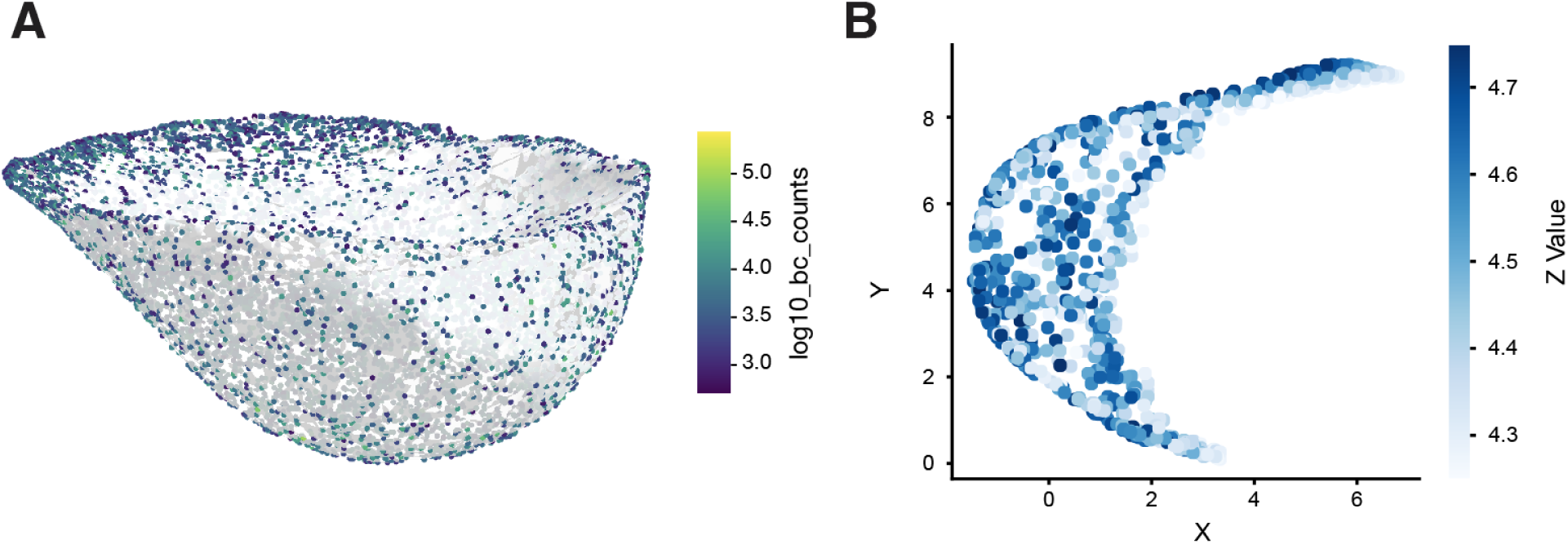
SCOPE recovers volumetric structure when bead recovery is not diffusion-limited. **(A)** 3D UMAP reconstruction of the settled-bead control, using n_neighbors=15 and 1,000 training epochs on the interaction matrix. Each barcode was normalized to 10,000 total counts per barcode and then log-transformed. Points are bead barcodes in 3D space overlaid on a concave hull (white) calculated by the Python package “alphashape” and colored by the log10 UMI count of each barcode. The reconstruction recapitulates the expected conical geometry of the tube bottom. **(B)** Z-axis cross-section through the reconstruction in panel **A**.

**Figure S9:**
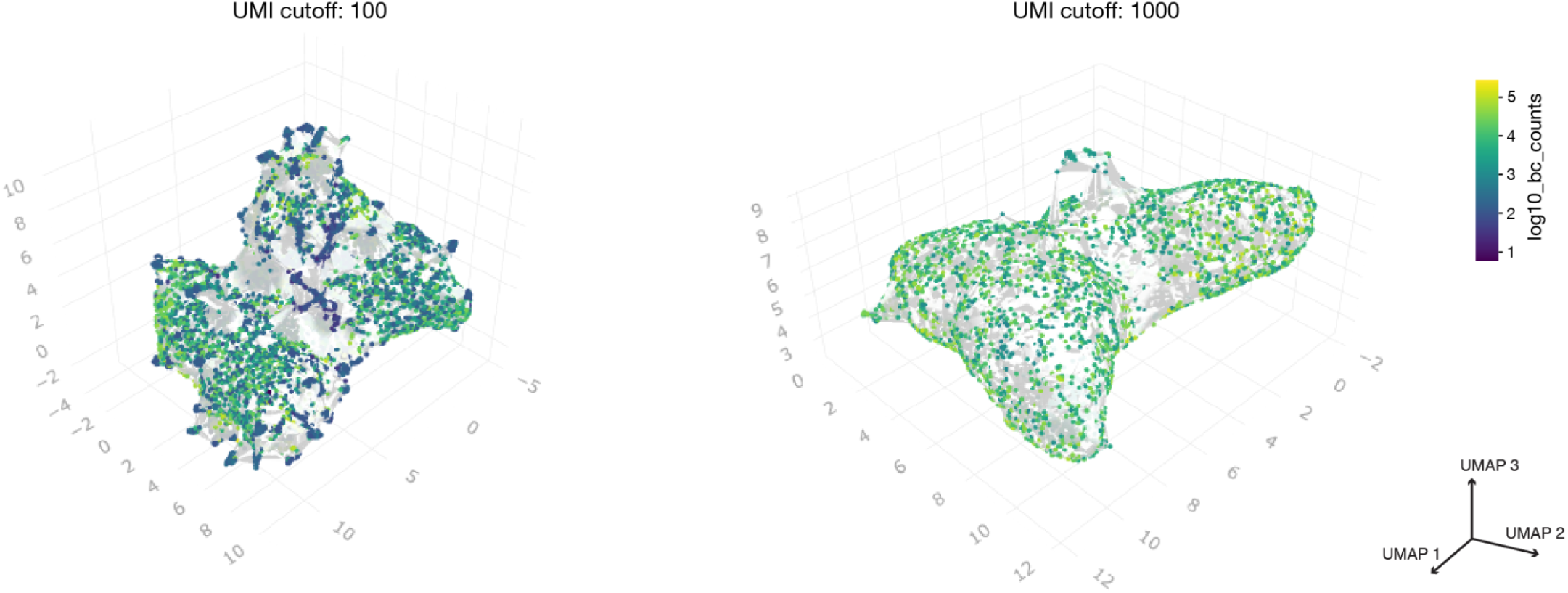
Lowering UMI cutoffs for barcode inclusion in 3D reconstructions leads to spurious barcode clusters. SCOPE sequencing libraries were filtered for barcodes with at least 100 UMIs (left) vs. 1,000 UMIs (right) and reconstructed in 3D using the UMAP algorithm (1,000 training epochs, *n_neighbors* = 15, *min_dist* = 0.4) on the normalized count matrix.

**Figure S10:**
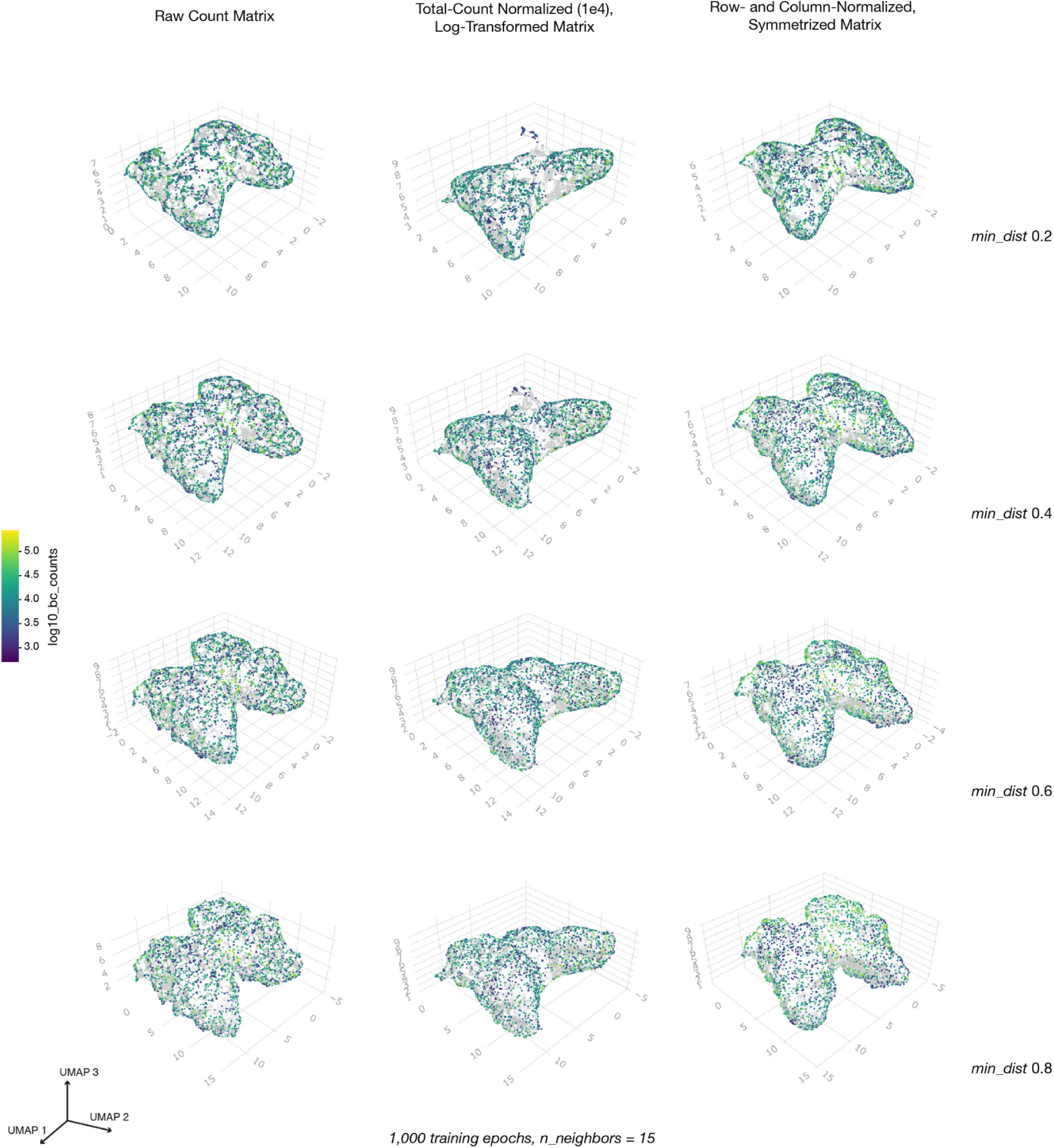
Representative results from a parameter grid search performed on the bead–bead interaction matrix derived from the butterfly-shaped hydrogel mold. The matrix was filtered to include barcodes with at least 1,000 UMIs and reconstructed using a fixed number of training epochs (n = 1,000) and nearest neighbors (n = 15). Columns show different normalization strategies: (left) raw count matrix; (middle) normalized count matrix scaled to 10,000 total counts per barcode and log-transformed; (right) matrix normalized by dividing each entry by its row and column sums, then symmetrized by averaging with its transpose. Rows correspond to varying *min_dist* values.

**Figure S11:**
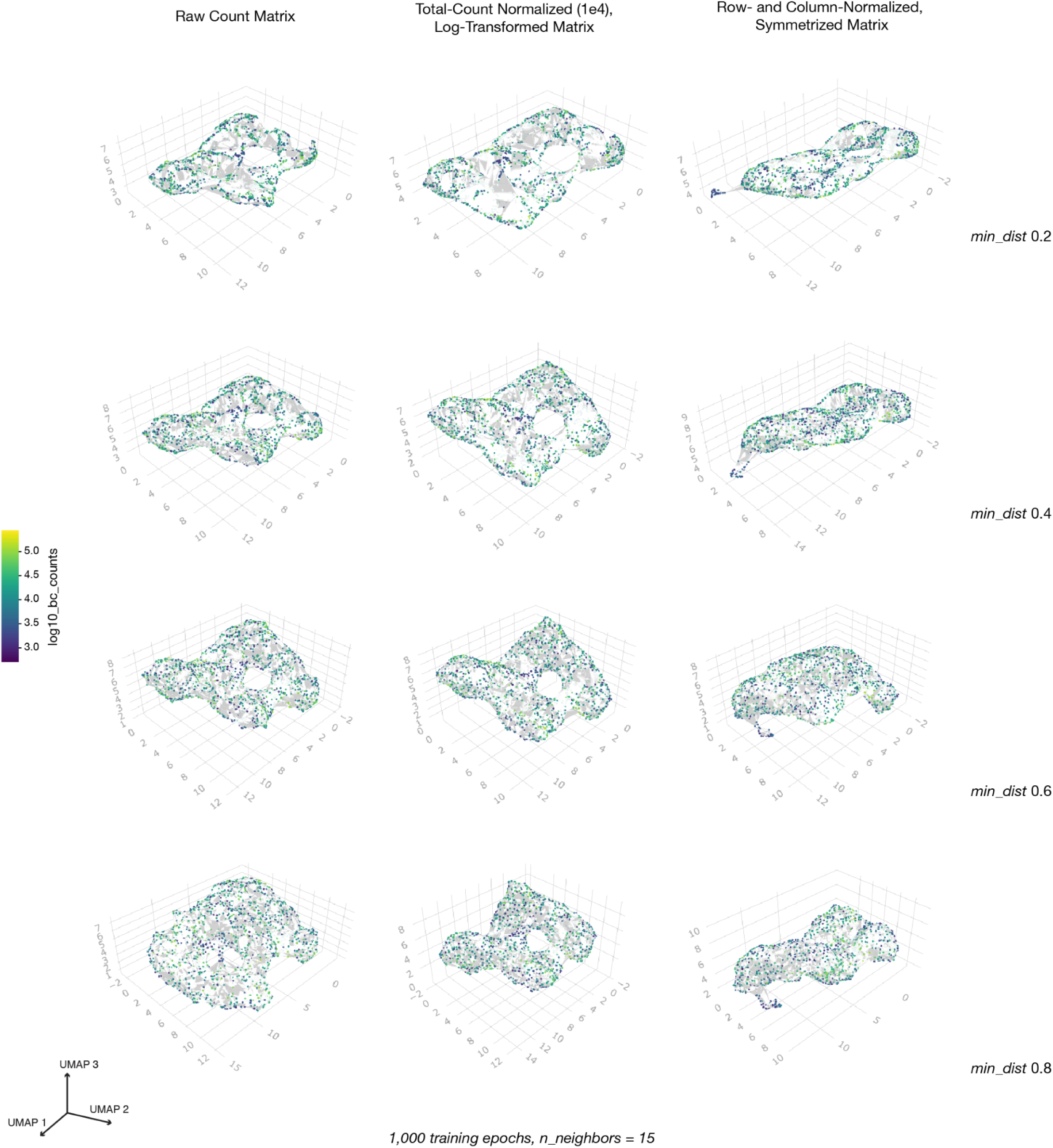
Representative results from a parameter grid search performed on the bead–bead interaction matrix derived from the block letter E-shaped hydrogel mold. The matrix was filtered to include barcodes with at least 1,000 UMIs and reconstructed using a fixed number of training epochs (n = 1,000) and nearest neighbors (n = 15). Columns show different normalization strategies: (left) raw count matrix; (middle) normalized count matrix scaled to 10,000 total counts per barcode and log-transformed; (right) matrix normalized by dividing each entry by its row and column sums, then symmetrized by averaging with its transpose. Rows correspond to varying *min_dist* values.

## Supplementary Materials

## Materials and Methods

### Molecular biology methods

#### Hydrogel bead fabrication

Hydrogel beads were produced through the use of a flow focusing microfluidic device to create water in oil emulsions (Zilionis et al., 2017). First, acrydite-modified oligos were designed and ordered from IDT with the TruSeqR1 (OLG_001: /5Acryd/TTTTTTT/ideoxyU/CTACACGACGCTCTTCCGATCT) and TruSeqR2 (OLG_002: /5Acryd/TTTTTTTTTTGTGACTGGAGTTCAGACGTGTGCTCTTCCGATCT) Illumina sequences. Acrydite oligos were stored at -20°C. The TruSeqR1 handle contained a deoxyuracil for USER enzyme-controlled cleavage off the bead. To make the polyacrylamide gel beads, a DNA and acrylamide mix (3% (v/v) acrylamide/bis solution (Sigma), 3% acrylamide solution (Sigma), 48 mM Tris–HCl pH 8.0, 0.25% (w/v) ammonium persulfate, 0.1X Tris-buffered saline–EDTA (TBSET: 10 mM Tris–HCL pH 8.0, 137 mM NaCl, 20 mM EDTA, 1.4 mM KCl, 0.1% (v/v) Triton-X 100), 50 µM of OLG001, and 50 µM of OLG002) was first prepared. The TEMED (Bio-Rad) catalyst was held out of the aqueous phase to prevent premature polymerization and added into the emulsion collection tube instead. The acrylamide mix was run through a droplet generator (Droplet Genomics) set up with a DG-DM-25 chip (Atrandi Biosciences) as the aqueous phase at 200 µL/hr, along with 2% RAN-008-FS v/v (Ran Biotechnologies) in HFE7500 (Oakwood Chemical) as the oil phase at 300 µL/hr. The resulting emulsions were allowed to polymerize overnight at room temperature. To break the emulsions, 150 uL of 1H,1H,2H,2H-Perfluorooctanol, 97% (Thermo Scientific Chemicals) was added for every 1 mL of beads produced. The bead solution was vortexed and centrifuged for 1 minute at 1000 x g and excess oil was removed from the polyacrylamide bead layer. Two more washes with a 1:1 ratio of hexane to TBSET were performed to further remove residual oil. Finally, multiple TBSET washes were performed until all the residual oil was removed.

#### Hydrogel bead barcoding and functionalization

Four consecutive rounds of splint barcodes were ligated together to build the bead barcodes. Eight plates of top and bottom oligos were ordered for the four barcode splints (**Table S1**). The following protocol closely followed the Delley & Abate protocol to make combinatorially barcoded hydrogel beads (Delley and Abate, 2021). Briefly, polyacrylamide beads were prepared for splint ligation by first annealing primer OLG_003 (**Table S2**) to the polyacrylamide-incorporated DNA stub on the bead. This step creates a four base pair overhang as a handle on which to ligate the first barcoded splint. Barcoded splints were ordered as two separate oligos to form the top and bottom portions of the splint and mixed at a 1:1 ratio. If the splint components were ordered without a 5’ phosphorylation modification, T4 PNK (NEB) was used to add phosphate groups to the top and bottom oligos before each round of ligation. For the first round of barcoding, beads were distributed across a 96 well plate containing barcoded splints and ligated with T4 ligase (NEB). The plate was placed in a thermomixer at 37°C for 2 hours, shaking at 1000 rpm to prevent beads settling at the bottom of the wells. The T4 ligase reaction was inactivated at 65°C for 20 minutes before the beads from each well were pooled together and mixed well. The barcoding and split-pooling steps were repeated for a total of four times to create a 96^4^ combinatorial barcode space.

Once the beads have been uniquely barcoded via splint ligation, a capping sequence was added to the end of the barcode to functionalize the bead oligo. Since the bead proximity reactions rely on creating 20 subtypes of beads that can interact with beads of another subtype but not with their own, the beads were divided into 20 pools for 20 separate functionalization reactions. Each functionalization reaction contains 1.5 parts of each of the 19 antisense capping oligos and 20 parts of the single sense capping oligo (**Table S1**; Sense_messenger_001–020, Antisense_messenger_001-020). For example, one of the reactions would contain 20 µL of Sense_messenger_001 and 1.5 µL each of Antisense_messenger_002 through Antisense_messenger_020 (Antisense_messenger_001 was left out). The entire mixture of 19 antisense to one sense oligos was then mixed with a poly-dT capping oligo (**Table S1**; Decoder_001) at a 1:1 ratio, and then ligated onto the fourth bead barcode splint with T4 ligase. Thus, both decoder and messenger species were loaded onto the bead barcodes and the beads were able to perform poly-A molecule capture and bead-bead proximity interactions, respectively. Finally, to get rid of incomplete DNA oligo stubs resulting from inefficiencies at any of the previous ligation steps, beads were treated with ExoI nuclease after annealing complementary oligos to all possible functional 3’ caps. Beads were then stored at 4°C for up to one year.

#### Casting densely packed monolayers of barcoded beads immobilized on glass slides

To immobilize barcoded beads in a dense monolayer, 15 µL of packed beads were first mixed with 2.75 µL of Bis-acrylamide 37.5:1 (Bio-Rad) to form a bead slurry. 0.55 µL of freshly prepared 5% (w/v) APS (Bio-Rad) was added to the bead slurry and the entire solution was pipetted as a large droplet onto a 25.4 mm x 76.2 mm microscope glass slide functionalized with 3-(Trimethoxysilyl)propyl methacrylate (Sigma-Aldrich) as described previously (Fu et al., 2022). A 22 mm x 22 mm coverslip was placed over the bead slurry and lightly pressed down to spread the beads into a single layer. These reagent volumes were scaled for casting larger arrays. The spreading of the beads was checked by eye under the microscope to ensure a densely packed monolayer under the area of the coverslip.

Once beads have been spread underneath the coverslip in the acrylamide mixture, we sought to polymerize the acrylamide directly onto the functionalized glass surface. As oxygen inhibits the polymerization reaction, a chamber was prepared using a one-gallon Ziploc bag filled with inert argon gas to flush out the oxygen or set in an anaerobic chamber (Coy Labs). To catalyze the polymerization of the encasing gel, 2% (v/v) TEMED solution was pipetted around the edges of the coverslip and the slide was placed in an anoxic chamber overnight. Once the gel had polymerized, the glass slide was incubated in a tray with enough water to cover the slide for at least 10 minutes at room temperature. After the encasing gel had been hydrated, a new razor blade was inserted under the edge of the coverslip and used to peel the coverslip away from the bead array. 1 mL of water was then used to rinse the bead monolayer for a total of five times, and the rinsed slide was placed into a tray with enough water to cover the monolayer for 10 minutes. These rinsing steps were for washing off unpolymerized acrylamide monomers in the encasing gel that inhibit subsequent PCR reactions. The bead monolayer was then dried uncovered in the fume hood and stored at 4°C until use.

#### Printing images on bead arrays with oligonucleotide inks

Thirteen poly-dA oligos (OLG_007 through OLG_019) were first reconstituted to 100 µM in IDTE pH 7.5 and then diluted to 1 µM in printing buffer (0.01% Tween-20 (v/v) and 0.5% (v/v) glycerol). Because the FAM-conjugated poly-dA oligo displayed weak fluorescent signal during imaging, DAPI was included into the printing buffer for this particular oligo so that those spots emitted fluorescence in the DAPI channel. Each oligo was loaded into a separate well of a 384 well-plate (Scienion CPG-5502-1) and spotted using the Scienion Sciflexarrayer S3 using a piezo dispense capillary 100 (PDC 100). A custom map was loaded for each oligo and prints were conducted serially with a spot-to-spot spacing of 100 µm with a pitch of 50 µm. After printing, the bead array was imaged using the Keyence BZ800 with a 4x objective. Tiled images were stitched and saved for image processing.

#### SCOPE reactions for messengers and decoders (circular and asymmetric arrays – *Fig. 3*)

The desired shape was made in the dried bead monolayer by scraping the excess beads off the glass slide with a clean razor blade. To constrain poly-dA hashing oligos to specific areas of the bead monolayer for capture, 0.2 µL of 0.1 µM poly-dA hashing oligo (**Table S2**; OLG_004 through OLG_006) was pipetted directly onto the bead monolayer and let dry in the fume hood. Next, a silicone chamber (Grace Bio-Labs, 20 mm diameter, 2.5 mm height) with an adhesive back was adhered to enclose the shaped bead monolayer. For the USER and reverse transcription reaction, a master mix of 1X CutSmart (NEB), 1X SuperScript IV RT Buffer (Thermofisher), 500 µM dNTPs (NEB), 67 U/mL of USER enzyme (NEB), 10000 U/mL of SuperScript RT Enzyme (Thermofisher) was prepared. For a 20 mm diameter chamber, 50 µL of USER and reverse transcription master mix was added to the center of the bead monolayer. An 18 mm diameter glass cover slip round was gently placed on top of the bead monolayer so that the master mix is spread evenly underneath the coverslip and covers the entire area of the monolayer.

The slide was then placed in a pre-warmed thermal cycler with an adaptor that can directly transfer heat evenly from the thermal cycler block to a glass slide. Care was taken to ensure that the lid of the thermal cycler does not press down on the slide with the adherent chamber, with PCR tubes placed at the edges of the thermal cycler block to keep the thermal cycler lid off the slide. The glass slide was then incubated at 37°C for 15 minutes and 55°C for 15 minutes for the USER and reverse transcriptase reactions, respectively. Once the USER and reverse transcription reaction were finished, the slide was taken out of the thermal cycler and a clean razor blade was used to carefully remove the coverslip on top of the bead monolayer. To collect the USER-cleaved barcodes from the bead monolayer, which form the sender-decoder and proximal poly-dA oligo chimera, 200 µL of water was added into the chamber and a P200 pipette was used to triturate the solution gently. The entire volume of solution inside the chamber was collected into a new 1.5 mL tube, which was then used for generating the library of poly-dA molecules that have been captured by a spatial bead barcode.

To collect bead interaction information, which is based on sequencing counts of sender-messenger and receiver-messenger chimeras, the bead monolayer in the chamber was washed three more times with 300 µL of water. After removal of the last wash, 100 µL of 0.1 M NaOH was added into the chamber, covering the bead monolayer. The solution was incubated at room temperature for 15 minutes to denature the sender-receiver chimeric molecules off of the beads. Then, the entire volume of the chamber was collected into a new 1.5 mL tube. 100 µL of 0.1 M Tris-HCl pH 8.0 was pipetted into the chamber and triturated gently several times with a P200 pipette to neutralize the previous NaOH wash and rinse the remaining denatured sender-receiver chimeric molecules from the bead monolayer. The entire volume was then collected from the chamber and added to the 1.5 mL tube with the previously collected NaOH wash.

A SPRI reaction was performed on the entire poly-dA collection volume to size select for the decoder library and eluted in 20 µL of water. On the entire volume of denatured sender-receiver messengers, a SPRI cleanup was performed and eluted in 20 µL of water. These two elutions were then prepared as Illumina sequencing libraries via an indexing PCR step to add the Illumina handles and sequencing indices.

#### Indexing PCRs for SCOPE libraries (circular and asymmetric arrays – *Fig. 3*)

A 30 µL indexing PCR reaction was prepared with 15 µL of NEBNext High-Fidelity 2X Master Mix (NEB), 10 µL of template DNA (SPRI purified from the previous section), and 4 µM each of forward and reverse indexing primers (**Table S2**). Messenger reactions used TruseqP5 (OLG_025) and Truseq P7 (OLG_026) indexing primers, while decoder reactions used NexteraP5 (OLG_028) and TruseqP7 (OLG_026) indexing primers. SYBR Green was added to track the number of PCR cycles before saturation and the PCR was performed with an annealing temperature of 69°C and an extension time of 45 seconds. The PCR reaction was stopped before the qPCR SYBR Green curve went past the exponential phase. A final SPRI cleanup was performed on the crude PCR product from the decoder-polyA molecule capture library, as well as a final SPRI cleanup of the crude PCR product from the sender-receiver messenger library. The concentration of the libraries was quantified using a High Sensitivity D1000 ScreenTape on a TapeStation 4200 system. Most sequencing runs were performed with P2 200 cycle kits on NextSeq 2000 (Illumina) with standard chemistry. Reads were paired-end spanning 105 base pairs for Read1, 105 base pairs for Read2, and 10 base pairs for both P5 and P7 indices. This read structure provides two times the sequencing coverage over the portion where annealing happens in the chimeric sender-receiver molecule. The same read structure can also accommodate the decoder-captured poly-dA molecule library, so that it can be sequenced on the same run as the messenger library. To increase sequence diversity, between 10 and 15% Phi-X was spiked into the sequencing library and runs were loaded at 850 pM.

#### SCOPE reactions for messengers and decoders (large eye exam array - *Fig. 4*)

Since the oligo spots that were printed onto the bead array show outwards diffusion as in **Fig. S7**, we sought to limit the diffusion of these printed oligos so that letter shapes with sharp boundaries can be recovered. After the oligos were printed on the bead array, three poly-dA oligos (OLG_020 through OLG_022) were diluted to 0.1 µM and a single oligo was used in consecutive washes of the array. Thus, we could soak up the available decoder molecules on the beads that were not in contact with the printed oligo spot through hybridization with these background poly-dA oligos.

Given the larger size of the bead array for the eye exam diagram reconstruction compared to the previous circular and asymmetric arrays, the reaction volumes were scaled accordingly based on the fold change in surface area of the arrays. A glass coverslip was placed on top of the bead array during the two-phase USER and reverse transcriptase reaction, as described for the circular and asymmetric arrays. To collect the supernatant containing the sender-decoder and poly-dA oligo chimeras, the array was washed with a low salt buffer (10 mM Tris-HCl pH 8, 10 mM NaCl, 3 mM MgCl2, 0.1% (v/v) Tween-20, 0.1% (v/v) NP-40). The messenger library was collected by incubating the array in 300 µL of 0.2 M NaOH at room temperature for 7 minutes and quenching with 50 µL of 1 M Tris-HCl pH 8. The collected decoder and messenger libraries were then SPRI purified as described for the circular and asymmetric arrays.

#### Indexing PCRs for SCOPE libraries (large eye exam array – *Fig. 4*)

For the decoder library with captured poly-dA hashes, a 200 µL indexing PCR reaction was prepared with 100 µL of NEBNext High-Fidelity 2X Master Mix (NEB), 90 µL of template DNA (SPRI purified from the previous section), and 0.2 µM each of NexteraP5 forward (OLG_028) and TruseqP7 reverse (OLG_026) indexing primers (**Table S2**). SYBR Green was added to track the number of PCR cycles before saturation and the PCR was performed with an annealing temperature of 60°C and an extension time of 30 seconds. The PCR reaction was stopped at 6 cycles. A 0.8X SPRI cleanup was performed on the crude PCR product from the decoder library to generate the final sequencing library.

For the messenger reactions containing bead proximity information, a 300 µL PCR reaction was prepared with 150 µL of NEBNext High-Fidelity 2X Master Mix (NEB), 100 µL of SPRI purified messenger products, and 0.2 µM each of TruseqP5 forward (OLG_023) and TruseqP7 reverse (OLG_024) primers (**Table S2**). These primers did not contain a sequencing index, which would be later added in the subsequent PCR reaction. SYBR Green was added to track the number of PCR cycles before saturation and the PCR was performed with an annealing temperature of 63°C and an extension time of 30 seconds. The PCR reaction was stopped at 8 cycles. Two consecutive 0.85X SPRI cleanups were performed. The SPRI purified template was then put into a 100 µL indexing PCR reaction with 50 µL of NEBNext High-Fidelity 2X Master Mix (NEB) and 0.2 µM each of TruseqP5 forward indexing (OLG_025) and TruseqP7 reverse indexing (OLG_026) primers (**Table S2**). Both libraries were sequenced on both the Illumina Nextseq 2000 and Novaseq as previously described for the circular and asymmetric arrays.

#### Fluorescence in situ hybridization

For FISH of fiduciary barcode probes, a similar procedure was followed where arrays were washed in 1 mL of 6X SSC three times before incubation with 1 µM of fluorescence-conjugated oligos (**Table S2**; OLG_029 and OLG_030). Primers were hybridized to the bead barcodes at 37°C for 30 minutes. After primer hybridization, unbound primers were removed and the array was washed 3 times with 1 mL of 1x PBS with 0.01% (v/v) Tween-20. The bead array was then imaged on the Keyence BZ-X800 microscope using EGFP and Cy5 filter sets from Chroma using a 10x objective (NA 0.45, Nikon).

#### Casting bead-packed 3D gel scaffolds

A dense slurry of 100 um DNA-barcoded beads was mixed into 4-6% final polyacrylamide concentration (15:1 acrylamide:crosslinker) and 0.7% APS. After coating the inside of the desired mold with 1-2 µL of TEMED, the bead slurry and encasing gel was pipetted into the mold and allowed to polymerize at room temperature for 20 minutes. Once polymerized, the gel matrix was released from the mold and soaked in 10 mL of water for 10 minutes. The polymerized gel was then washed with 1 mL of water for a total of three times. Gels were stored in water until ready for use on the same day.

#### SCOPE reactions for bead-embedded gel matrices (3D reconstructions – *Fig. 5*)

The polymerized bead-embedded gel was then transferred to a 2 mL-Eppendorf tube. A 150 µL master mix was prepared with 75 µL of NEBNext High-Fidelity 2X Master Mix (NEB), 8 µL of USER (NEB), and 1X rCutSmart (NEB) and transferred to the sample. Depending on the size of the polymerized gel, a larger volume tube and higher volume of master mix may be required to submerge the gel in the master mix. The tube was then incubated at 37°C for 30 minutes and slowly ramped up to 72°C over the course of another 30 minutes.

To collect bead interaction information, which is based on sequencing counts of sender-messenger and receiver-messenger chimeras, the polymerized gel was washed three times with 1 mL of water. After removal of the last wash, 150 µL of 0.1 M NaOH was added into the tube, submerging the bead-embedded gel object. The solution was incubated at room temperature for 10 minutes to denature the sender-receiver chimeric molecules off of the beads. Then, the entire volume of the tube was collected into a new 1.5 mL tube. 150 µL of 0.1 M Tris-HCl pH 8.0 was then pipetted into the tube and triturated gently several times with a P200 pipette to neutralize the previous NaOH wash and rinse the remaining denatured sender-receiver chimeric molecules from the beads. The entire volume was then collected from the chamber and added to the 1.5 mL tube with the previously collected NaOH wash. For larger 3D matrices, a larger volume of denaturing and quenching solution may be required to submerge the object.

On the entire volume of denatured sender-receiver messengers, a 1.0 X SPRI cleanup was performed and eluted in 20 µL of water. The elution was then prepared for Illumina sequencing via an indexing PCR step to add the Illumina handles and sequencing indices (same indexing PCR reactions as the ones for 2D SCOPE reactions).

### Computational methods

#### Fastq read processing for bead-bead interaction matrix

From the fastq file of sequencing the messenger library, the four 10-bp sub-barcodes of the combinatorial spatial barcode were extracted from Read1 and error corrected to known barcodes through the following process. In our final bead barcode, known 4-bp “scars” exist between each of the sub-barcodes from ligating the barcoded splints together. Additionally, the set of 96 sub-barcodes at the first position contain extensions between 0 and 3-bp, which need to be determined to extract the rest of the sub-barcodes. To do so, each pair of sequencing reads was first mapped to scar sequences to determine the extension length of the first of the four sub-barcodes (between 0 and 3-bp), as well as the positions of each sub-barcode. Next, the four extracted sub-barcodes were mapped to true sub-barcode sequences. The annealed portion between the sense and antisense ends of two bead barcodes were also mapped to known messenger sequences.

For error correction of the sequences, certain numbers of substitutions were allowed to account for sequencing errors via calculating the Levenshtein distances between them. The Levenshtein distance between two sequences is the minimum number of single-base edits (insertions, deletions or substitutions) required to change one sequence into the other. For scars mapping, we tried different extension lengths and took the extension length that minimizes the Levenshtein distance and then allowed the Levenshtein distance to be lower than 2. For barcode mapping, we allowed each of the 4 barcodes to have a Levenshtein distance lower than 2. For mapping of the annealed region between the two barcodes, we corrected sequences up to 2 Levenshtein distance away from known messenger sequences.

#### Fastq read processing for poly-dA oligo capture library

From the fastq file of sequencing the decoder library, the four 10-bp sub-barcodes of the combinatorial spatial barcode were extracted from Read1 and error corrected to known barcodes following the same process as the one for the messenger library. The poly-dA oligo’s barcode was extracted from the first ten base pairs of Read2, and the number of UMIs from the Read1 spatial barcode indicated how many poly-dA oligos were captured.

#### Quantifying bead-bead Interactions

After error-correcting the sequencing reads for the messenger library, we performed 3 steps of filtration to determine real barcodes in our dataset. We first filtered the reads by collapsing interactions between the same pairs of UMIs. Next, we filtered the barcodes by the number of times they appear, keeping barcodes that appear at least “x” number of times, which was determined by where the steepest drop-off occurred in a knee plot of UMI counts. Finally, we filtered for barcodes that appear in both reads sent from beads and reads from receiving beads. We then encoded each bead barcode into a unique integer, and counted the number of interactions between every pair of barcodes. These steps result in an asymmetric interaction count matrix where the rows are barcodes being sent from beads and the columns are barcodes being received by beads. This interaction count matrix was then stored in the sparse matrix format.

#### Doublet detection to remove sets of beads that share the same DNA barcode

Since large numbers of beads were present in the arrays, a non-negligible number of beads may be doublets, i.e. beads that do not have a unique DNA barcode. The theoretical number of doublets in an array is a function of the number of possible barcodes (N) and the number of beads in the array (n):

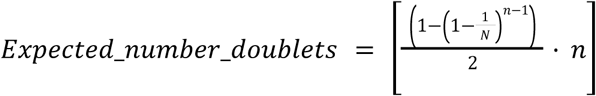

We performed clustering-based doublet detection to remove such beads. We first built an unweighted graph with the interaction count matrix: if two beads had non-zero interaction counts, there was an unweighted edge between the nodes representing these two beads in the graph. Next, we iterated through each node in the graph and performed Leiden clustering to find communities among its neighbors. The threshold we set for a bead being a doublet was as follows: the neighborhood of the bead should form more than 1 cluster and the largest cluster must be less than 4 times the size of the second largest cluster. Since the number of doublets detected depends on the selected clustering resolution and varies across datasets, for each dataset we performed a grid search on the resolution parameter. We selected the clustering resolution that identified a number of doublets closest to the theoretical value.

#### Custom simulation of diffusing messenger oligos from beads arrayed in hexagonal lattice

We developed a simulator to guide development of inference methods. To develop the simulator, we did an empirical investigation to understand the decay of bead-bead interaction across distance, and landed on an inverse-square model (**Fig. 2H**). We also used a simple binomial model of bead compatibility. To do a full simulation, we combined the inverse-square weight function with a sample dataframe giving joint row and column read counts to get a per-bead sender and receiver read count, then used multinomial sampling to get the desired number of reads. This ensured that the joint distribution of read counts in the simulation matched those from real data.

#### Mapping from sender/receiver interactions to pairwise bead distances

Sequencing of the sender-receiver reaction products yields a bead-bead interaction count matrix. This matrix represents the counts of all observed interactions involving an oligo diffusing through space from one bead to another. Thus, it is in essence a “similarity” matrix; higher counts indicate beads that are closer together in space. However, it is not a symmetric measure, as the sending and receiving counts for each pair of beads are represented separately. This similarity matrix is highly sparse, as interactions for most pairs of beads that are not sufficiently close are not observed. Since most methods for reconstructing position data require Euclidean distance matrices as input, we sought to devise a way to map from our similarity matrix, i.e. the measured counts of sender/receiver interactions, to a distance matrix.

Using our custom simulation, we can generate such interaction count matrices alongside simulated bead positions, arranged in a hexagonal lattice. The simulation is parametrized with the per-bead messenger oligo count distribution from the empirical data. After normalizing this count matrix by the total number of molecules sent and received by each bead and averaging the matrix with its transpose, we arrive at a symmetric similarity matrix. We then use a random forest model to perform regression between the normalized pairwise bead-bead similarities for each pair of simulated beads and the corresponding pairwise Euclidean distances from the simulation. We then use this regression function, trained on a simulation, to map our observed, normalized similarity matrix from an experiment to an inferred pairwise distance matrix.

However, this distance matrix has several flaws, chiefly that all the “zeros” in the similarity matrix have been mapped to a single distance value, as per the nature of a regression function. Therefore, our distance matrix is only an approximation of the true pairwise distances between the beads, since the diffusion process that we observe in the form of our interaction matrix only measures discrete molecular binding events. Thus, an important property is that our distance matrix is most accurate at local distances. This precludes the use of classical methods such as Multi-Dimensional Scaling to compute the inferred bead positioning.

#### Clustering beads and computing global reconstruction

Rather, we opted to use Uniform Manifold Approximation Projection (UMAP) to compute the inferred bead positioning. Most often used as a manifold learning algorithm for dimensionality reduction and visualization, UMAP algorithms can also accept as input a pairwise precomputed k-nearest neighbor distance matrix. Although UMAPs are usually said to introduce significant error in dimensionality reduction, our problem is fundamentally a 2-dimension-to-2-dimension mapping problem, and thus avoids the reconstruction error issue. The superior performance of UMAP at larger scales according to our simulated results informed the choice of UMAP over t-Distributed Stochastic Neighbor embedding (**Fig. S3A**).

We apply three pre-processing steps to the data. First, we identify “doublet beads” according to our method described above, and remove them. Using the normalized similarity matrix as an adjacency matrix, we can construct a graph such that beads are represented by vertices/nodes and edges between them are weighted by the normalized similarity. We then apply a published method for pruning “short circuit edges” in the bead-bead interaction graph that confound the reconstruction (Kloosterman et al., 2024). Finally, we compute the “k-core” of the bead-bead interaction graph after doublet beads and spurious edges have been filtered out-that is, we compute the largest subgraph such that all nodes have a minimum degree of “k”, which is set to 100 (using an implementation in the NetworkX package). This is done to ensure that only beads with sufficient information for spatial mapping purposes are kept.

This filtered bead-bead interaction graph is then clustered using the Leiden community detection algorithm to group beads into clusters of at most 2500 beads. We convert the pairwise bead-bead similarity matrix to a pairwise distance matrix using the method described above. However, since we would like to run UMAP with a nearest neighbors value higher than 100 (the minimum degree in the network), we need to partially impute the “missing” pairwise distance values for which there were no observed bead-bead interactions. To do this, we predict the pairwise distance value for 0 counts using our regression function, and within each cluster of beads, we impute any missing pairwise distances with this value. In this way, we can then compute a k-nearest neighbor pairwise distance matrix with k=250, and use this precomputed sparse distance matrix as input to UMAP. We initialize the UMAP using the PAGA algorithm(Wolf et al., 2019). PAGA provides a force-directed layout visualizing relationships between the bead clusters, in a manner that accurately represents the shape of the underlying manifold. We use this to provide initial estimates of the centroids of each cluster, and then initialize the beads in each of these clusters as being randomly distributed around these centroids according to a 2D Gaussian distribution. We first compute an “initial UMAP” using this as its initialization and using fixed hyperparameters.

We perform a grid search over the “min_dist” and “repulsion_strength” UMAP hyperparameters in order to refine the solution. We define two metrics for hyperparameter selection; the inferred solution should be contiguous (since we expect one unified sheet of beads, not broken up into “islands”), and the beads in the solution should be as evenly spaced as possible. To assess the first criterion, we represent each putative reconstruction (for each hyperparameter value) as a binary image such that each pixel, each spanning an equal area of the reconstruction point cloud, is colored black if it has any points within it. This image is then “eroded” using the scikit-image package to remove artifacts, followed by image segmentation using the Chan-Vese algorithm. All solutions with more than one large object detected are discarded.

Finally, from the set of such valid solutions, we select the solution that has the most evenly spaced points by first computing a 2D histogram of the point cloud to quantify the density map of the inferred bead positions. We then calculate a chi-squared test statistic between this distribution of per-bin bead densities and the uniform distribution, and select the solution that has the lowest test statistic as being closest to a uniform density of beads. In this way, we are able to select the most optimal UMAP result over the hyperparameters in the grid search.

#### Downsampling experiment and evaluation of t-SNE vs UMAP

In order to determine how the reconstruction accuracy varies with the sequencing depth of the sender/receiver library, we designed a synthetic downsampling experiment. We simulated a 40,000 bead array using the framework above, parametrized using the counts of the asymmetric array experiment. We then downsampled the total counts of the simulated bead-bead interaction matrix to various levels ranging from 1% to 100% of the total counts. For each chosen downsampling percentage, we attempted to reconstruct the spatial positions of the beads and align them to the true known positions of the beads according to the simulation.

This registration was done in the following way. In a real-world experiment, if more information is provided on bead positions through bead segmentation on brightfield or fluorescence microscopy images, we could then align the inferred positions to the true positions. The alignment can be done by matching the positions of “fiducial” beads, identified through hybridization to an oligo probe complementary to a partial bead barcode sequence. However, since the precise one-to-one matching of fiducial beads is not known, we apply an algorithm for the linear sum assignment problem, or the minimum weight matching problem for bipartite graphs (implemented in scipy). In essence, the set of inferred positions for the fiducial beads can be represented as one partition of vertices, while the true positions represent the other partition. The weights of the edges between these partitions of vertices are defined by the pairwise distances between their respective positions. The linear transformation that aligns the fiducial bead positions is learned through gradient descent, recomputing the bead point matching at every iteration and minimizing the mean Euclidean error of the matched points. The linear transformation is parameterized according to 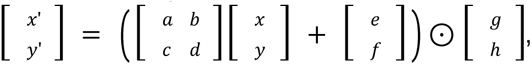 for a total of 8 free parameters. After an initial alignment of the fiducials, the point matching is then computed for all points. An optimal linear transformation to align all the points is then performed and the point matching is recomputed.

The final point matching is then used to determine the final positions of the beads. Instead of using the inferred positions of the beads, we use the positions of their inferred match, in effect “snapping” the inferred positions onto the experimental lattice. In this way, we eliminate much of the positioning error as long as the matches are correct. The result from these steps thus represents the more refined result that can be obtained using simple brightfield microscopy and segmentation, and was applied to results from the simulation in order to quantify reconstruction error (**Fig. S3**). We repeated this process 10 times for each downsampling percentage and computed the mean Euclidean error of the inferred bead positions after alignment (**Fig. S3**). On the x-axis, we plot the mean messenger UMI counts per bead after downsampling.

In order to evaluate the accuracy of t-SNE vs UMAP at various scales, we simulated rectangular hexagonal lattice bead arrays at various different sizes and computed the t-SNE and UMAP reconstructions on this bead-bead interaction data without breaking up the graphs into clusters. The reconstructions were then registered and point matched to the ground truth as above, and the mean Euclidean error was calculated. The median and interquartile range of 5 trials were plotted for each simulated array size.

#### Reconstructing the letters in the Snellen eye chart

For the Snellen diagram experiment, there were a few differences in how the reconstruction was performed due to the scale of the experiment. First, instead of using the full “k-core” algorithm to remove beads with low degree, we simply removed all beads with less than a degree of “k” in one step. Although this does not ensure that the resulting graph will have a minimum degree of “k”, this was done for memory reasons and yields roughly similar results. Second, in order to minimize observed distortions in the reconstruction, we kept the remainder of the pipeline the same but computed the initial UMAP using 3 dimensions, rather than 2. This 3D UMAP was then reduced to 2D to produce the final result in a second UMAP step, computed using a kNN with a Euclidean distance metric on the 3D result (k=50), yielding our final result (**Fig 4B**). We were unable to perform the hyperparameter gridsearch, also due to memory constraints. We then visualized individual letters by cropping the corresponding regions in the reconstruction (**Fig 4C**). We created a binned image of the large letter “E” by tiling the reconstruction point cloud with equally sized squares, and summed up the squared counts of the captured DAPI oligo to visualize the banding pattern (**Fig. 4D**). Horizontal and vertical line plots were computed using the values in the bins that the lines intersected at each axis coordinate.

#### Segmentation of imaged fiducials and alignment with reconstruction

To validate the accuracy of our local reconstructions, we sought to determine whether the results agreed with a partial ground truth obtained through optical sequencing. To this end, we stained the SCOPE array with a Cy5 channel (OLG_029) and FITC channel FISH probe (OLG_030) which hybridized to two distinct groups of barcodes found only on a small subset of beads. The SCOPE array was imaged using a Keyence microscope at 10x (NA 0.45, Nikon) using the EGFP and Cy5 filter sets from Chroma. We then applied segmentation and point registration on the acquisitions in the following manner. First, image channels from the acquisition were merged into a single gray scale image by summing all pixel intensities. Next, we subtracted the background intensity and split the single-channel image into 36 equally sized tiles to optimize shape segmentation by Meta AI’s Segment Anything Model (v1). This deep learning model was used to generate masks and centroids of FITC beads using the model’s default parameters. These centroids were then merged together to generate a final point cloud representing beads hybridized by the FITC probe. The positions of the beads with barcodes that match the FITC probe targets were registered using the methods described in the global reconstruction section above. The optimal linear transformation that was learned was then applied to the Cy5 probes as secondary validation.

## Supplemental Text

**<u>Figs. S1 to S9</u>**

Legends inline with text

**<u>Tables S1 to S2</u>**

**Table S1. Oligonucleotide sequences to generate barcoded and functionally capped SCOPE beads.** The full bead barcode consists of four sub-barcodes that are 10 base pairs each. Two 96 well plates of oligonucleotide sequences are provided within each sheet, named “BC1”, “BC2”, “BC3”, and “BC4”. The two plates consist of the top and bottom sequences of a barcoded splint with ligation overhangs. Once fully barcoded with four rounds of splint ligation, beads are capped with one species of a decoder sequence for capturing polyA-molecules, and twenty species of messenger sequences for capturing barcode-barcode interactions. The twenty pairs of complementary messenger sequences used in SCOPE, as well as the decoder sequence, are provided in the “Functional caps” sheet.

**Table S2. Oligonucleotide sequences for SCOPE bead array experiments.** Poly-dA oligonucleotides used for the oligo-painted image, sequencing primers to generate the messenger and decoder libraries, and FISH primers are included.

## Notes

https://github.com/SrivatsanLab/SCOPE

https://github.com/matsengrp/sci-space-v2

### Summary of Updates

Edited discussion, updated references and new supplemental data on volumetric diffusion of oligos.

